# Caloric restriction promotes beta cell longevity and delays aging and senescence by enhancing cell identity and homeostasis mechanisms

**DOI:** 10.1101/2023.08.23.554369

**Authors:** Cristiane dos Santos, Shristi Shrestha, Matthew Cottam, Guy Perkins, Varda Lev-Ram, Birbickram Roy, Christopher Acree, Keun-Young Kim, Thomas Deerinck, Melanie Cutler, Danielle Dean, Jean Philippe Cartailler, Patrick E. MacDonald, Martin Hetzer, Mark Ellisman, Rafael Arrojo e Drigo

## Abstract

Caloric restriction (CR) extends organismal lifespan and health span by improving glucose homeostasis mechanisms. How CR affects organellar structure and function of pancreatic beta cells over the lifetime of the animal remains unknown. Here, we used single nucleus transcriptomics to show that CR increases the expression of genes for beta cell identity, protein processing, and organelle homeostasis. Gene regulatory network analysis link this transcriptional phenotype to transcription factors involved in beta cell identity (Mafa) and homeostasis (Atf6). Imaging metabolomics further demonstrates that CR beta cells are more energetically competent. In fact, high-resolution light and electron microscopy indicates that CR reduces beta cell mitophagy and increases mitochondria mass, increasing mitochondrial ATP generation. Finally, we show that long-term CR delays the onset of beta cell aging and senescence to promote longevity by reducing beta cell turnover. Therefore, CR could be a feasible approach to preserve compromised beta cells during aging and diabetes.

## Introduction

Aging is associated with loss of normal cell and tissue function, which are linked to degenerative and metabolic diseases such as Alzheimer’s and diabetes during old age ^1^. Tissues composed of largely post-mitotic cells are expected to be significantly impacted by aging because most cells in these tissues can be remarkably long-lived and as old as the organism itself ^2–4^. Because of their longevity, long-lived cells (LLCs) are under an immense pressure to maintain their normal cell function for long time periods to sustain organ function. However, how LLCs achieve and maintain structure and function during adulthood, and whether aging-associated deficits in LLC function in old organisms can be reversed to restore normal tissue function remains largely unknown.

Using a combination of in vivo stable isotope labelling of mammals (SILAM), correlated and high-resolution electron and multi-isotope mass spectrometry (called MIMS-EM), and protein mass spectrometry pipelines we have identified the tissue distribution of LLCs and of long-lived protein (LLP) complexes in post-mitotic cells ^4–8^. These studies identified that different endocrine cell types in the mouse pancreas are LLCs, including up to 60% of all insulin-secreting beta cells ^4,9,10^. Beta cells secrete insulin in response to increases in blood glucose levels to sustain normal glucose homeostasis for an entire lifetime^11^. Loss of beta cell insulin secretion can occur during normal aging and is the cause of type 2 diabetes (T2D); this phenotype is linked to compromised expression of transcription factors (TFs) that maintain beta cell identity ^12–14^, accumulation of islet fibrosis, inflammation, and ER stress ^12,15^, reduced KATP channel conductance^16^ and loss of coordinated beta cell calcium dynamics ^17^, impaired beta cell autophagy and accumulation of beta cell DNA damage that triggers beta cell senescence ^12,14,18,19^. Aging-and T2D-associated impairment of beta cell function are also characterized by an increase in transcriptional noise and re-organization of beta cell gene regulatory networks (GRNs) that normally support beta cell structure-function, including the beta cell ER stress response ^12,20^.

Caloric restriction (CR) and CR-mimicking approaches (e.g., time restricted feeding (TRF)) can prolong organismal longevity and delay aging from yeast to non-human primates ^21–26^. These beneficial effects are associated with improved glucose homeostasis due to prolonged fasting and enhanced peripheral insulin sensitivity, enhanced insulin signaling, lower adiposity, enhanced mitochondrial homeostasis and lower ATP generation, increased autophagy, enhanced protein homeostasis and reduced ER stress and inflammation ^21–29^. In the pancreas, 20-to-40% CR or CR achieved via TRF are linked to lower islet cell mass in lean mice^21^, whereas in pre-diabetic mice CR restores normal beta cell secretory function, identity, and preserves beta cell mass^30–33^ in a process dependent on activation of beta cell autophagy via *Beclin-2*^34^. In patients with a recent T2D diagnosis, the efficacy of extreme CR (average of 835 kcal/day or >50% CR based on a 2,000kcal/day diet) to reverse T2D depends on the capacity of beta cells to recover from previous exposure to a T2D metabolic state^35^. However, the mechanisms underlying the adaptation of beta cells to CR and whether CR can delay beta cell aging remains largely unknown.

We investigated these questions by exposing adult mice to mild CR (i.e., 20% restriction) for up to 12 months and applied *in vivo and in vitro* metabolic phenotyping of beta cell function followed by single cell multiomics and multi-modal high resolution microscopy pipelines (electron, light, mass spectrometry, and isotope microscopy) to investigate how CR modulates beta cell heterogeneity and longevity. Our data reveals that CR reduces the demand for beta cell insulin release necessary to sustain euglycemia by increasing peripheral insulin sensitivity. This causes beta cells to undergo a significant reorganization of their transcriptional architecture that promotes a largely post-mitotic and long-lived beta cells with enhanced mitochondrial structure-function while mitigating the onset of aging and senescence signatures. Therefore, our results provide a molecular footprint of how CR promotes beta cell function and longevity to delay aging and that could explain the beneficial effects of CR to the treatment of T2D in humans.

## Results

### Calorie restriction (CR) modulates beta cell function in vivo

We exposed 8-week-old FVB or mice to 20% CR for 2-months, and quantified in vivo glucose homeostasis mechanisms (Figure 1A). As expected, CR mice consumed 80% of the total daily kilocalories (kcal) consumed by control mice maintained in an ad libitum (AL) diet (Figure S1A). After 2 months, CR mice did not experience significant body weight gain due to a reduction in fat mass and adiposity, and maintained lean body mass (Figure 1B, Figure S1B). In contrast, mice exposed to high-fat diet (HFD) for 2 months consumed more calories and became obese (Figure 1B-C, Figure S1A).

**Figure 1.**
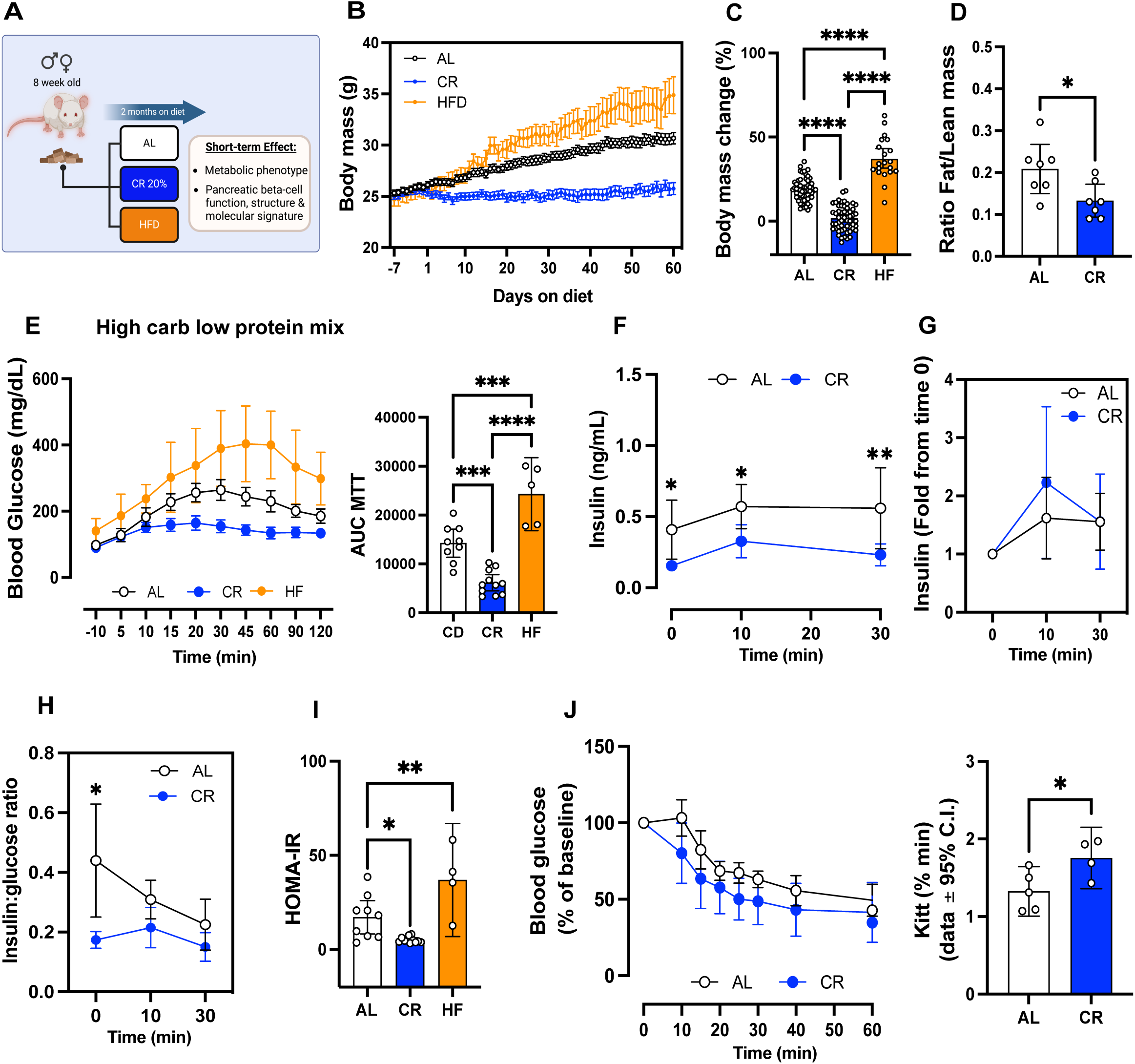
Short-term of caloric restriction (CR) improves glucose homeostasis in mice. **(A)** Schematic diagram of mice subjected to ad-libitum (AL), 20% CR, or HFD for 2 months starting at 8 weeks of age. **(B)** Mouse body mass over 2 months on diet. **(C)** Body mass change after 2 months on diet. **(D)** Ratio between fat mass and lean mass after 2 months on diet. **(E)** Blood glucose levels during the meal tolerance test (MTT) after 2 months on diet and respective areaunder-curve (AUC) measurements. **(F)** Insulin levels during the MTT and **(G)** the respective fold change from baseline values. **(H)** Ratio between the insulin and glucose values obtained during the MTT. **(I)** HOMA-IR calculated from fasting glucose and insulin values after 2 months on diet. **(J)** Blood glucose values obtained during an intra-peritoneal insulin tolerance test (ipITT) and the respective decay of the glucose rate per minute (kITT). All data are presented as mean ± 95% confidence intervals (C.l.) The asterisks mean * *p*< 0.05, ** *p*< 0.01, *** *p*< 0.001 and **** *p*< 0.0001 using one or two-way ANOVA with Tukey’s post hoc test or Sídák’s multiple comparisons test. Unpaired Student t-test was applied for two groups comparation in **(D, J).** In **(B-C),** n=20-75 male mice per diet group; **(D)** n=7 male mice per diet group; **(E)** n=5-11 male mice per diet group; **(F-I)** n=9-11 male mice per diet group; **(J)** n=5 male mice per diet group.

To quantify the impact of CR on beta cell insulin release and glucose homeostasis *in vivo*, we performed an oral mixed-meal tolerance test (MTT) using a high-carbohydrate nutrient shake (Ensure, ∼2% of the daily kcal consumption, see methods for details). This revealed that CR mice have improved glucose tolerance compared to AL mice, whereas HFD mice were glucose intolerant (Figure 1E), however fasting glucose levels were not different from AL mice (CR (n=26) 105.4±14.81 mg/dL versus AL (n=24) 114.0±19.34 mg/dL, p=0.0834). Serum insulin measurements before and during the MTT revealed that CR beta cells secrete ∼50% less insulin than beta cells in AL mice (Figure 1F). Similar experiments using higher doses of glucose (2g glucose/kg of body weight) confirmed these results (Figure S1C-E); importantly, the relative capacity of CR beta cells for stimulated-insulin secretion was similar to that of AL beta cells (Figure 1G, Figure S1E), which suggests that CR does not impair beta cell stimulated insulin secretion mechanisms. In fact, CR beta cells secrete approximately half the amount of insulin to sustain normoglycemia than AL mice due to increased insulin sensitivity (Figure 1H-J). The lower levels of circulating insulin in CR male mice were not due to changes in activity of the hepatic insulin degradation enzyme (IDE, Figure S1F) or to changes in beta cell mass (Figure S1G).

Similar experiments in 8-week-old C57/BL6 male confirmed these results (Figure S2A-I). In addition, no changes in fasting glucagon levels (AL (n=11): 2.473±1.473 versus CR (n=12) 2.896±1.240 pM) or in alpha cell mass were observed (Figure S1G). In contrast, experiments in 8-week-old female mice demonstrated that CR induces similar changes to body weight mass and composition (Figure S3A-D), while it fails to improve glucose tolerance or insulin sensitivity or alter *in vivo* beta cell function (Figure S2E-K). These results could be explained by the fact that female mice already have significantly higher peripheral insulin sensitivity than male mice and CR does not further improve insulin sensitivity in relation to AL-fed mice (Figure S3L).

Next, to measure beta cell insulin release in vitro, we performed dynamic glucose-stimulated insulin secretion assays (GSIS) using isolated islets from male mice exposed to AL, CR, or HFD for 2 months to measure beta cell insulin release *in vitro*. Surprisingly, no significant differences in basal and/or glucose-stimulated insulin release or islet insulin content were observed between diet groups (Figure S1H-I). However, CR beta cells displayed reduced insulin release when challenged with high glucose in combination with the phosphodiesterase inhibitor 3-isobu-tyl-1-methylxanthine (IBMX) (Figure S1H). These results suggested that CR beta cells could have lower glucose-dependent cAMP generation, which is required for amplification of glucose-dependent insulin secretion and incretin signaling^36,37^. To test this hypothesis, we repeated the *in vivo* MTT in AL or CR mice pre-treated with the GLP1 receptor agonist Liraglutide (400ug/kg body weight, 2 hours before the start of the assay). Pre-treatment with Liraglutide improved glucose tolerance in both groups and sustained higher serum insulin levels after 30 minutes in AL but not in CR mice (Figure S1J-M). Therefore, this data indicates that CR beta cells might have reduced GLP1R signaling *in vivo* in conditions of meal-induced and transitory hyperglycemia.

Together, these results suggest that CR improves peripheral insulin sensitivity and glucose tolerance in a sex-dependent manner, which decreases the demand for beta cell insulin secretion necessary to maintain normal glucose levels in male mice without significantly affecting glucose-dependent insulin release mechanisms or alpha cell function.

### CR alters beta cell epigenetic and transcriptional heterogeneity

Given that 2 months of CR was sufficient to trigger significant changes to glucose homeostasis and beta cell insulin secretion *in vivo*, we applied single nucleus (sn) transposase-accessible chromatin with sequencing (ATAC)-and mRNA sequencing (called snMultiome-seq, 10x Genomics) to investigate the transcriptional architecture underlying the adaptation of beta cells to CR or HFD after 2 months on diet.

Single nuclei were isolated from frozen islet pellets from mice exposed to AL, CR, or HFD diet for 2 months (Figure 2A). We analyzed a total of 16,555 high-quality islet nuclei (Figure 2B, see methods) to quantify chromatin accessibility and coverage of our sequencing assay and identify major islet cell types using the relative expression and chromatin accessibility of well-established islet cell marker genes (Figure 2C-F, Figure S4A-F)). To determine the effect of CR on overall beta cell gene expression, we performed a pseudo-bulk differential gene expression analysis comparing AL, CR, and HFD beta cell transcriptomes. Relative to AL beta cells, CR beta cells show up regulation of several beta cell identity genes, including both insulin genes (*Ins1*, *Ins2*), amylin (*Iapp*), the insulin processing enzyme *Pcsk1n*, the glucose-6 phosphatase enzyme *G6pc2*, the beta cell transcription factor (TF) *Nkx6-1*, and down-regulation of the incretin receptor *Gipr* (*Glp1r* was not detected) and of the carbohydrate-responsive transcriptional regulator *Mlxipl* (also known as *ChREBP*) (Figure 2G, Supplementary Table 1).

**Figure 2.**
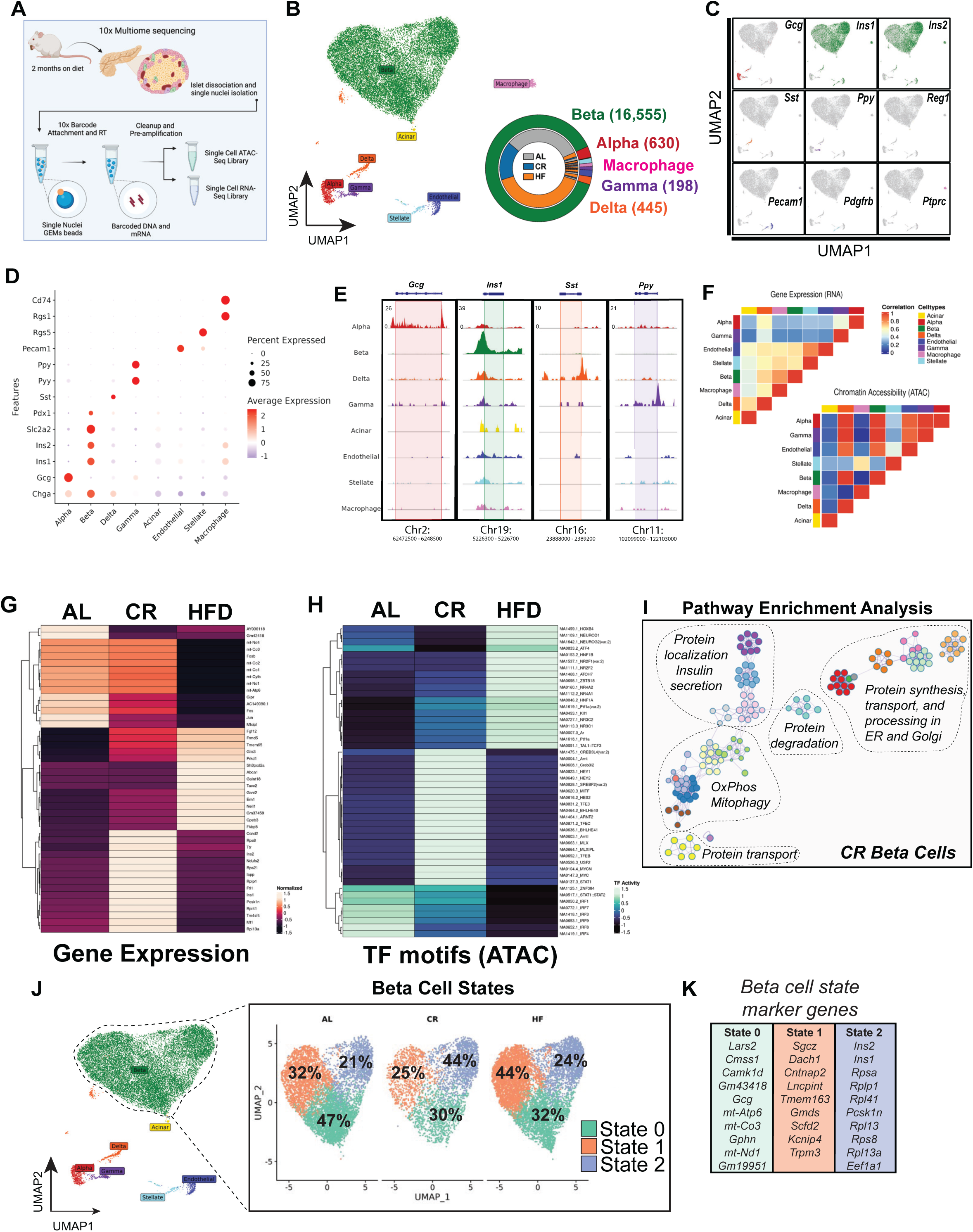
Single cell multiome sequencing of islets reveals diet-specific changes to betacell heterogeneity. **(A)** Schematic representation of the workflow used for single nuclei multiome sequencing (ATAC + RNA) of pancreatic islets isolated from AL, CR and HFD mice after 2 months on diet. **(B)** Uniform Manifold Approximation and Projection (UMAP) of the integrated transcrip-tome and chromatin dataset. Data from a total of n=18,741 islet cells from n=2 mice per diet group. Different cell types are indicated by various colors and labels. Inset, donut plot with the total number of cells in each cell cluster. **(C)** UMAPs showing the expression of cell marker genes. **(D)** Dot plot with relative expression of marker genes in alpha, beta, delta, gamma, acinar, endothelial, stellate and macrophage cell clusters. **(E)** Genome tracks showing ATAC peaks for hormone markers: Gcg (alpha cell), Ins1 (beta cell), Sst (delta cell) and Ppy (gamma cell). **(F)** Heatmap showing the correlation between gene expression (RNA-seq) and chromatin accessibility (ATAC-seq) across the identified cell types. **(G)** Heatmap with the top differentially expressed genes (RNA-seq) and **(H)** top ATAC-seq peaks in transcriptional factors (TF) motifs in beta-cells from AL, CR, and HFD mice. **(I)** Annotated node map with pathway enrichment analysis of differentially expressed genes and TF motifs in beta-cells from CR mice. Analysis was performed using Metascape with an FDR < 0.05. **(J)** UMAP projection of beta-cells and identified subclusters from AL, CR and HFD mice. Beta-cell populations were defined according to distinct transcriptional states. **(K)** Top marker genes from each beta-cell state.

CR had a limited effect on beta cell chromatin accessibility (n=143 differentially modulated chromatin with p.val < 0.05), where most changes were associated with genes containing TF motifs mapped to circadian rhythm-(*Arntl*/*Bmal*, *Dbp*, *Tef*, and *Hlf*) and nutrient-dependent regulators of autophagy (*Tfeb* and *Tfe3*) (Figure 2H, Supplementary Table 2). Recent studies have shown that beta cells with higher levels of *Arntl/Bmal* are more functional and with increased expression of beta cell identity markers such as *Pdx1* ^38,39^. Immunohistochemistry (IHC) and confocal microscopy of islets from CR mice confirmed that ∼80% of all CR beta cells have higher levels of *Arntl/Bmal* and Pdx1 *in situ* after 2 months on diet (Figure S4G). Pathway enrichment analysis of differentially expressed genes and enriched chromatin regions in CR beta cells revealed the up regulation of pathways linked to beta cell identity and function, oxidative phosphorylation (OxPhos), mitophagy, protein processing, degradation, and transport through the endoplasmic reticulum (ER) and Golgi apparatus (Figure 2I). In contrast, HFD beta cells showed reduced expression of genes associated with beta cell function (i.e., *Fos*, *Fosb*, *Jun*, and the incretin receptor *Gipr)*, and increased expression of stress-and diabetes-associated genes (*Fkbp5* and *Glis3*, respectively) (Figure 2G-I).

Next, we identified three different beta cell transcriptional states (beta cell states 0, 1, and 2) and their marker genes and chromatin accessibility profiles (Figure 2J-K, Figure S4H-J). This analysis shows that CR caused a shift in the transcriptional heterogeneity of beta cells, where CR mouse islets had ∼2x more beta cells in state 2 (versus AL and HFD islets (Figure 2J)). Beta cells in this transcriptional state have higher expression of beta cell identity and function (*Ins1, Ins2, Mafa*), oxidative and anaerobic carbohydrate metabolism (*Pkm, Idh2, Acly*) and mitochondrial structure and function (Cytochrome C genes (*Cox4-to-8)*, *Vdac1*, *Tomm20*, and *Tomm70a*), autophagy (*Lamp1, Cltc, p62/Sqstm1, Ulk1, Ctsb*), lipid metabolism (*Hadh*), amino acid transport and metabolism (*Slc7a2*, *Slc38a2*), protein folding and response to ER stress (*Hspa5, Atf6, Creb3l2*) and DNA damage (*Nme1*) genes (Figure 2K, Figure S5A-E and S8C-F, Supplementary Table 3). Moreover, state 2 beta cells had lower expression of the rate-limiting enzyme glucokinase (*Gck*) and increased expression of glucose-6-phosphatase catalytic subunit 2 (*G6pc2)*, which could suggest that these cells have lower glycolytic flux and increased glucose excursion ^40^. However, this does not appear to be the case since CR beta cells have appropriate glucose-stimulated insulin release dynamics (Figure 1, Figure S1).

Together, these results indicate that CR triggers significant transcriptional and chromatin accessibility changes in most beta cells that enhance cell identity, nutrient metabolism, circadian rhythm regulation, and ER, mitochondria, and protein homeostasis mechanisms.

### CR reprograms beta cell gene regulatory networks (GRNs)

Beta cell development, maturation, and aging processes are characterized by modulation of gene regulatory networks (GRNs) that control beta cell gene expression patterns and heterogeneity ^12,41,42^. Given that CR has limited effects on beta cell chromatin accessibility (Figure 2, Figure S4-5), we wanted to test if the transcriptional phenotype of CR beta cells was largely due to re-organization of beta cell GRNs. To test this hypothesis, we applied the TF inference tool SCENIC ^43,44^ to scan the transcriptional signatures of individual beta cell to infer and assign TF-gene linkages using a known TF motif database to create cell type-specific GRNs ^12,43,44^.

We applied SCENIC to our single beta cell snMultiome-seq dataset (n=16,555 cells in total) from AL, CR, and HFD mice after 2 months on diet. We identified a total of 242 TF enriched in all beta cells, including beta cell identity TFs *Pdx1*, *Nkx-6.1, Mafa, and Foxa2a* (Figure 3A, Supplementary Table 4). Next, we determined the overall degree of correlation in the inferred activity of each TF to identify modules of co-regulated TFs (as we have previously shown ^12^). This approach revealed the expected and positive correlation in the activation of beta cell TFs (*Pdx1*, *Mafa*, and *Foxa2*, (Figure 3B)^45^. In addition, we identified distinct sets of co-regulated beta cell TFs, including the insulin-sensitive *Foxo1*, the ER homeostasis and stress response *Atf6*, the beta cell identity *Nkx6.1* and the carbohydrate-responsive *Mlxipl*/*ChREBP*, and two large groups of TFs associated with control of gene transcription mechanisms and cell development (*Sox9*, *Klf3/4*, *Runx1-3*) (Figure 3B).

**Figure 3.**
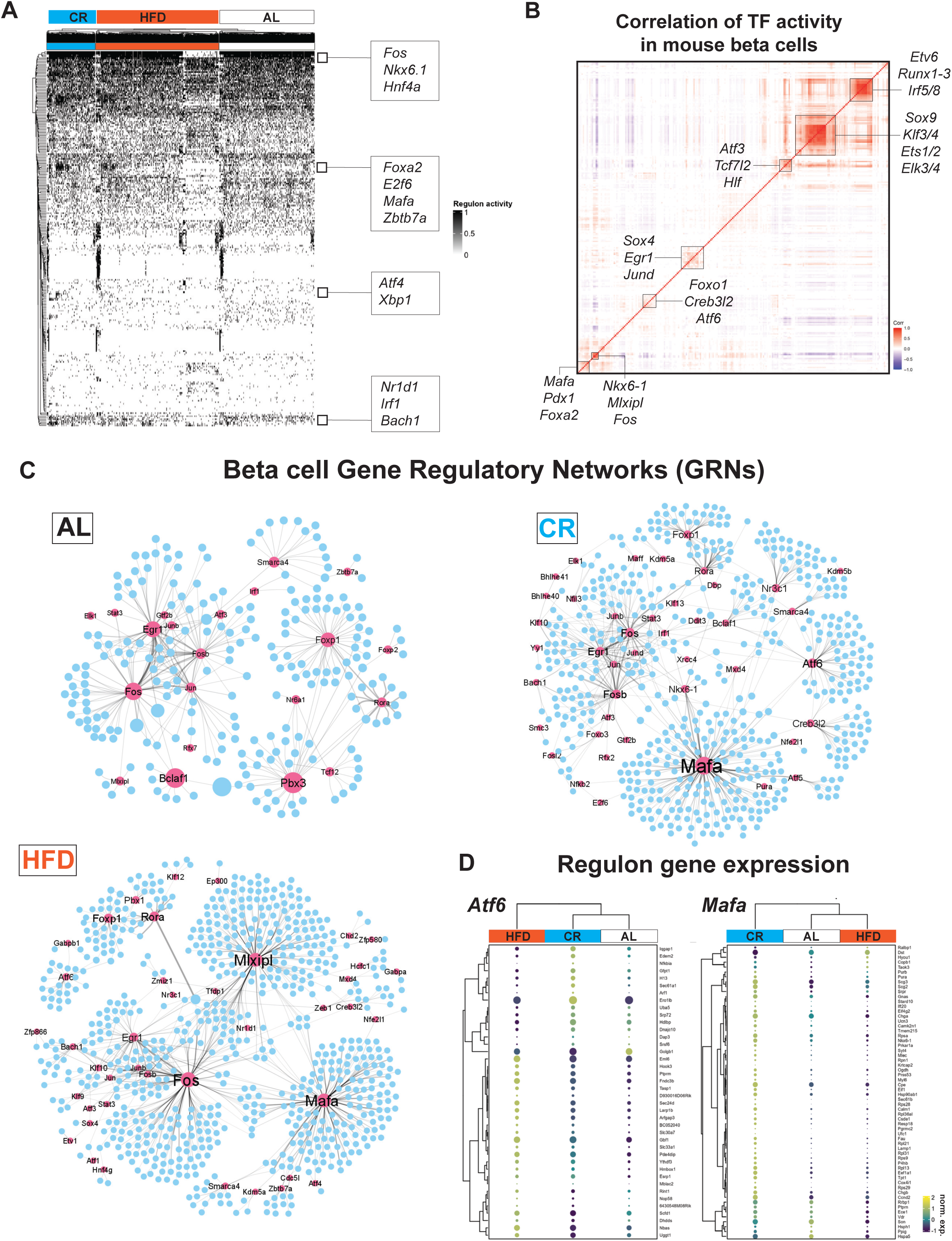
CR reprograms beta-cell gene regulatory networks (GRNs). **(A)** SCENIC heatmap with hierarchical clustering analysis of TF activity in beta-cells from AL, CR and HFD mice after 2 months on diet. TFs identified as “ON” are shown in black, while TFs identified as “OFF” are in white. **(B)** Pearson correlation matrix of n=272 TF identified in all mouse beta cells (all diet groups together). Boxes highlight clusters of TFs with high degree of correlation. **(C)** Gene regulatory networks (GRNs) formed by TFs identified using SCENIC in beta-cells from AL, CR and HFD mice. TFs are shown as pink nodes, while target protein-coding genes are shown in blue. Node size represents the “betweenness centrality” measurements that report in the influence of a given TF within a network. **(D)** Dot plot and hierarchical clustering showing the gene expression levels of target genes associated to Atf6 and Mafa GRNs from AL, CR and HFD beta cells. Dot plot scale shows the relative expression level and percentage of cells expressing a target gene.

Next, we used SCENIC to reconstruct diet-specific beta cell GRNs and to identify modules of TFs with similar activity patterns (Figure 4C, Figure S6A-D) and found that CR and HFD are associated with a significant reprograming of beta cell GRNs and TF activities (Figure 3C). Compared to AL beta cells, the CR beta cell GRN was characterized by increased activation and/or coordinated activity of TFs involved in beta cell identity and function (*Mafa, Nkx6-1, Foxo1*), protein folding and ER homeostasis (*Creb3l2, Atf6*), transcriptional regulation of mitochondrial genes (Gabpa/Nrf2 and Gabpb1), NAD metabolism (Sirt6), and glucocorticoid signaling (*Nr3c1*) genes, which coincides with increased gene expression of some of these markers in CR beta cells (Figures 2-3, and Figure S6D). In contrast, the GRN in HFD beta cells had increased activity of the glucose-responsive TF *Mlxipl*/*ChREBP*, which is required for glucose-stimulated beta cell proliferation and diabetes ^46,47^, and of the beta cell identity TF *Mafa* (Figure 3C). Notably, the network of genes targeted by *Mafa* in CR and HFD beta cells is not similar. For example, in CR beta cells *Mafa* is linked to autophagy and ER homeostasis genes *Beclin1* and *Calmodulin1,* respectively, whereas in HFD beta cells *Mafa* is linked to master regulators of beta cell proliferation (*Ccnd1*, *Cdc42*) and ER stress response (*Atf4*) (Figure S6E, Supplementary Tables 5-6). Similar results were observed for the beta cell proliferation TF *Foxp1*, and the protein homeostasis TF *Crebl2* (Figure S6E). Importantly, this beta cell TF target “reprogramming” was reflected in the gene expression pattern of genes associated with *Mafa* and *Atf6*. Here, these TFs in CR beta cells were linked to the increased expression of several beta cell identity and protein homeostasis genes (Figure 2-3), including the beta cell-specific ER chaperone *Ero1lb*, the alpha and beta subunits of the ER translocation channel Sec61 (*Sec61a1, Sec61b*), the regulator of intra-golgi transport *Arf1*, the autophagy marker *Lamp1*, and beta cell identity and function genes *Ucn3*, *Chga*, and *Nkx6-1* (Figure 3D).

Therefore, CR promotes beta cell identity and homeostasis by upregulating the expression of clusters of genes associated with enhanced beta cell maturation, homeostasis, and function. Moreover, this transcriptional program is likely driven by the increased transcriptional activity of *Mafa*/*Nkx6.1* and of other TFs shown to be involved in the maintenance of beta cell protein homeostasis and survival mechanisms (e.g., Atf6 ^48^).

**Figure 4.**
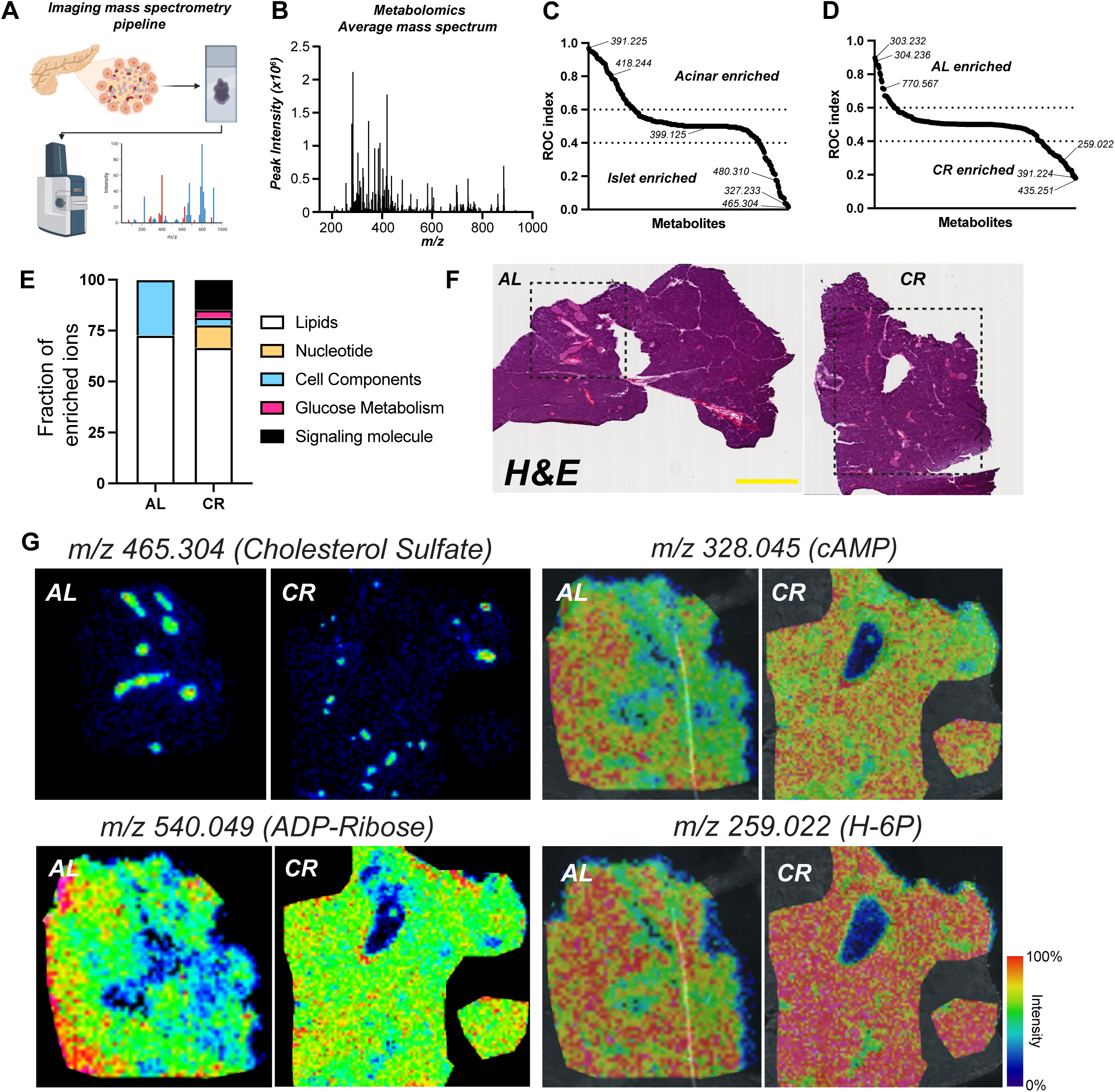
CR beta cells are metabolically fit. **(A)** Schematic diagram of imaging mass spectrometry (MALDI-MS) approach used to measure metabolite abundance in AL and CR pancreases after 2 months on diet. **(B)** Average mass to power (*m/z*) spectra of all samples combined. **(C)** Receiver Operating Characteristic (ROC) analysis of ions enriched in acinar versus islet regions. **(D)** ROC analysis of ions enriched in AL versus CR islet regions. **(E)** Fraction of MALDI-MS metabolites identified using the HMDB and their respective molecular class and distribution in AL or CR datasets. **(F)** Representative hematoxylin and eosin (H&E) staining of AL/CR pancreases prepared for MALDI-MS imaging. **(G)** Representative *m/z* images for select metabolites identified in our ROC analyzes. In **(F)**, scale bar = 3mm.

### Imaging mass spectrometry defines the spatial metabolomic landscape of CR islets

Our data indicates that CR changes beta cell function via modulation of beta cell identity, nutrient metabolism, and organelle homeostasis pathways (Figures 1-3). Next, we decided to investigate the metabolic profile of *in situ* islets using imaging mass spectrometry (called MALDI-MS) to quantify the relative abundance and spatial profile of metabolites in islets from AL and CR mice after 2 months on diet. We chose this approach to minimize changes in cell metabolism that could occur during the islet isolation process. Briefly, fresh mouse pancreases were snap-frozen using a liquid nitrogen bath and placed in histology cassettes prior to MALDI-MS metabolite imaging and data analysis (see methods for details) (Figure 4A). Consecutive sections were stained using Hematoxylin and Eosin (H&E) to pinpoint the spatial location of each islet (Figure 6A).

Our MALDI-MS metabolite imaging of the mouse pancreas identified a total of n=290 ions in AL and CR samples combined (Figure 4B); next, to identify metabolites enriched in islets versus acinar compartments we performed a Receiver Operating Characteristic (ROC) univariate analysis followed by metabolite database query. This approach revealed the identity of ∼50% of all ions detected and those significantly enriched in acinar (n=66 ions) or islet regions (n=42 ions) (Figure 6C, Supp Table 7). Acinar-enriched metabolites include the phospholipid cyclic phosphatidic acid (*m/z:* 391.22486, CPA 16:0) and nucleotide cytidine 2’ 3’-cyclic phosphate (*m/z:* 304.03402, cCMP), which is involved in the pyrimidine metabolism pathway. In contrast, islet-enriched metabolites were largely classified as lipids, including cholesterol sulfate (m/z: 465.30443) and phospholipids (CPA 18:0, or different types of phosphatidic acids (PA) (Figure 4C and G, Supplementary Table 7-9).

Next, we applied a similar analysis to identify metabolites enriched in AL or CR islets; this identified n=28 ions enriched in AL islets versus n=52 ions enriched in CR islets (Figure 4D). Our database search revealed that CR islet-enriched metabolites were classified between lipids, nucleotides, glucose metabolism, and cell components and signaling pathways (Figure 4E). These include the pyrimidine nucleotide signaling molecules uridine 2’,3’-cyclic phosphate (cUMP) and CMP, hexose 6-phosphate (glucose-6-phosphate or fructose-6-phosphate, m/z: 259.02236, G/F6P), cAMP (m/z: 328.04528), ADP (m/z: 408.0112), and cyclic ADP-ribose (m/z: 540.05334) (Figure 4D-G, Supp Table 7-8). Synthesis of ADP-ribose requires metabolism of nicotinamide adenine dinucleotide (NAD), which is increased in other cells exposed to CR and linked to glucose metabolism ^49,50^; importantly, ADP-ribosylation is required for proper DNA damage repair^51^ and this pathway is increased in CR beta cells (Figure S5A-E).

Together, this data shows that CR promotes changes to the overall islet lipid composition and increases the levels of molecules involved in beta cell glucose metabolism and insulin release (i.e., G/F6P, cAMP), and indicates that CR islets could have increased DNA damage response via ADP-ribosylation.

### CR delays beta cell aging

Aging beta cells accumulate ER stress^12^ and DNA damage, both of which are thought to contribute towards beta cell senescence and loss of function during old age ^18,19,52^. We hypothesized CR would prevent the onset of beta cell aging and senescence signatures and to test this hypothesis, we exposed 8-week-old male mice to CR for 12 months (Figure 5A). Long-term CR in mice is associated with reduced body weight due to lower fat content, arrested pancreas growth and a trend towards lower beta cell mass, and overall lower meal-stimulated beta cell insulin release due to increased peripheral insulin sensitivity (Figure 5B-H, Figures S1G and S7A). To investigate the effect of CR on beta cell aging signatures *in situ* and at the single cell level, we performed pancreas immunohistochemistry (IHC) followed by confocal microscopy of mouse pancreases from animals maintained on AL or CR for 2-to 12-months using the DNA damage response marker *53bp1*. Here, we observed that CR beta cells have reduced accumulation of 53bp1, thus indicating that these cells accumulate less DNA damage over time (Figure 5I) and that short term CR (i.e., 2 months) is already sufficient to induce benefits in DNA damage response, as indicated by our single cell transcriptomics analysis (Figure 2). To investigate additional markers of beta cell senescence, we performed IHC of 12-month-old mouse beta cells using the nuclear lamina marker *LmnB1* (which is lost in an accelerated model of mouse beta cell senescence ^53^) and mRNA Fluorescent In Situ Hybridization of *p16/Cdkn2a* and *p21/Cdkn1a* genes. We found that CR beta cells have significantly higher *LmnB1* levels and reduced *in situ* expression of *p16/Cdkn2a* and *p21/Cdkn1a* (Figure 5J-K, Figure S7B-C). Notably, most beta cells analyzed had p16 (96% AL vs 94% CR, p>0.05) or p21 (61% AL vs 59% CR, p>0.05) transcripts in nuclear and cytoplasmic compartments; however, no significant correlation in the expression of these markers was found at the single cell level (Figure S7D).

**Figure 5.**
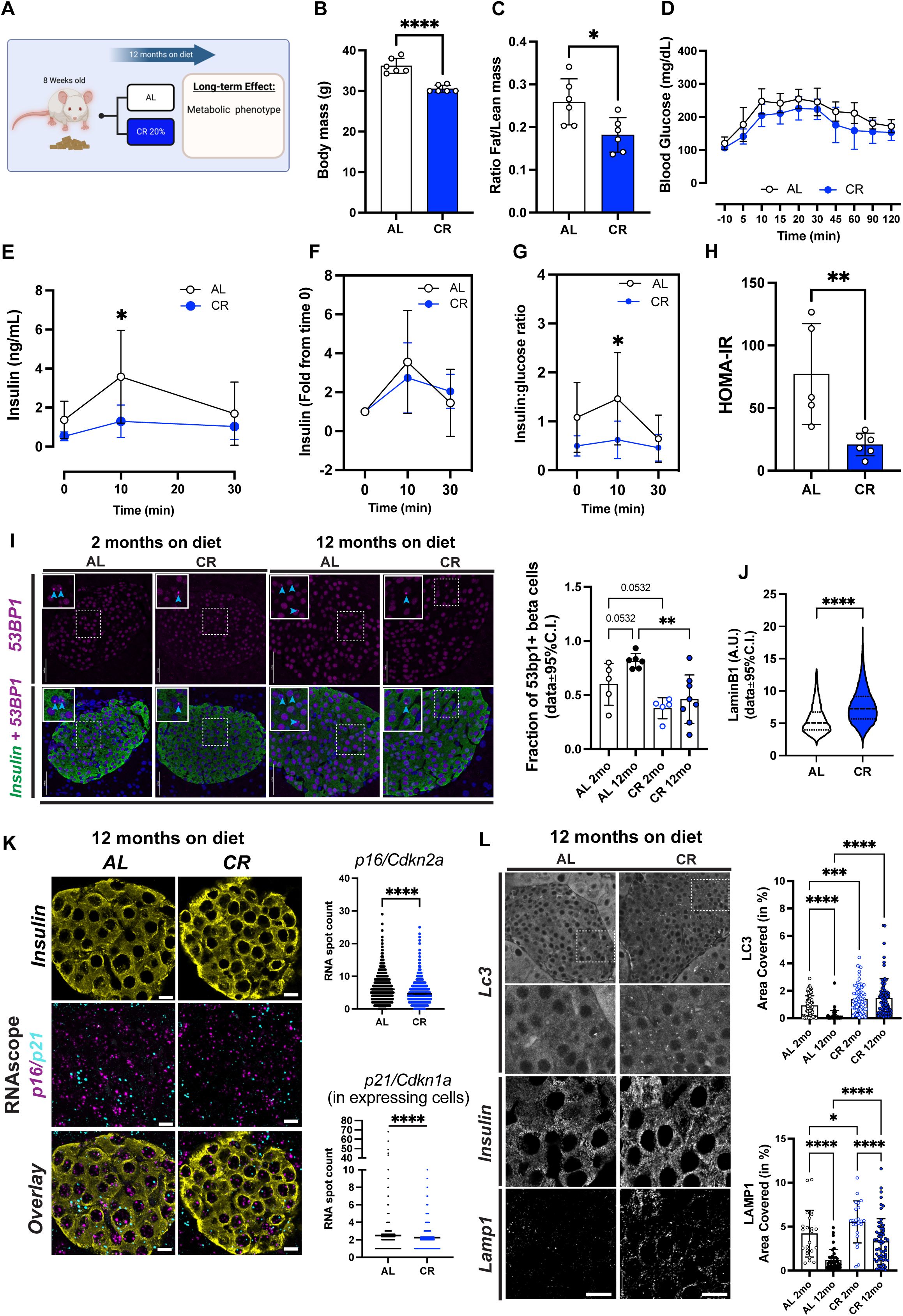
Long-term CR delays beta cell aging signatures. **(A)** Schematic diagram of mice subjected to AL or 20% CR for 12 months starting at the age of 8 weeks. **(B)** Body mass after 12 months on diet. **(C)** Ratio between fat mass and lean mass after 12 months on diet. **(D)** Blood glucose levels during the meal tolerance test (MTT) after 12 months on diet. **(E)** Blood insulin levels during the MTT and **(F)** the respective fold change from baseline values. **(G)** Ratio between the insulin and glucose values obtained during the MTT. **(H)** HOMA-IR calculated from fasting glucose and insulin values after 12 months on diet. **(I)** Representative images of pancreatic sections from AL and CR male mice after 2 months or 12 months on diet stained with 53BP1 and insulin. Right, quantification of DNA damage by 53BP1+ beta-cells. Each dot represents the average from each mouse. A total of 127-200 islets per experimental group were analyzed. Data are normalized by percentage of 53BP1 positive beta-cells per total beta-cell number. **(J)** Quantification of beta cell nuclear levels of Lamin B1 in pancreatic islets from AL and CR mice after 12 months on diet. **(K)** Representative images of pancreatic sections from AL and CR male mice after 12 months on diet stained with Cdkn2a and Cdkn1a mRNA probes, and insulin. Right, the quantification of incidence of these markers per beta-cell. **(L)** Representative images of pancreatic sections from AL and CR mice after 12 months on diet stained with Lc3I-II or Lamp1. Right, quantification of beta cell area occupied by Lc3I-II or Lamp1. For all panels, data are presented as mean ± 95% confidence intervals (C.l.). The asterisks mean * *p*< 0.05, ** *p*< 0.01, *** *p*< 0.001 and **** *p*< 0.0001 using two-way ANOVA with Sídák’s multiple comparisons test or unpaired Student t-test **(B, C, H, J, K).** In **(B-I)**, n=6 male mice per diet group.

Together, this data indicates that long-term exposure to CR maintains an insulin sensitive state that lowers the demand for beta cell function and insulin release, which reduces accumulation of aging-associated DNA damage and beta cell senescence markers.

### CR enhances autophagy and beta cell mitochondria function

Previous data indicates that autophagy is activated by intermitted fasting in HFD-fed mice to maintain beta cell mass and function ^34^, and our transcriptomics analysis identified up regulation of autophagy genes (*Beclin-1, Lamp1*) and transcriptional regulators of mitochondrial genes (*Gabpa/Gabpb1*) (Figure 2-3). First, we investigated the impact of short-and long-term CR on beta cell autophagy using the autophagosome marker *Lc3I-II* and the lysosome marker *Lamp1*, the later which is up regulated in CR beta cells (Figure 2). CR significantly increased both *Lc3* and *Lamp1* vesicle density in beta cells after 2 or 12 months on diet (Figure 5K-L, Figure S7E) and prevented the loss of *Lc3I-II* and *Lamp1* positive vesicles during aging of AL mice (Figure 5K-L). Autophagy is regulated by the nutrient sensor and cell growth regulator mammalian target of rapamycin (mTOR) that acts (in part) by phosphorylating Ser 242/244 of the ribosomal protein S6 (rpS6) to inhibit autophagy^54^. We measured rpS6 phosphorylation (p-rpS6) in beta cells from AL or CR mice *in situ* and found that CR led to significant down-regulation of p-rpS6 in beta cells (Figure S7F-G). This data indicates that mTOR activity is suppressed in beta cells, which would allow beta cells to increase autophagy during CR.

Autophagy is activated during conditions of nutrient deprivation is important for cell survival by supplying metabolites generated from the degradation of organelles and macromolecular complexes in lysosomal compartments ^55^. To determine the role and potential targets of beta cell autophagy during CR, we used deconvolution-assisted confocal colocalization microscopy of mouse pancreases stained with antibodies against lysosomes (*Lamp 1),* insulin granules (*Ins*), and/or mitochondria (succinate dehydrogenase isoform A (*Sdha*)). We found no differences in the co-localization between *Ins* and *Lamp1* across diet groups, which means that CR-induced autophagy does not target mature insulin granules (Figure S8A). Importantly, CR beta cells have lower levels of *Sdha*-*Lamp1* colocalization (Figure 6A), suggesting that beta cell mitophagy is reduced in these cells – which could be explained by the lower expression of the main mitophagy adaptor protein Parkin (*Prkn*) in CR beta cells (Figure S8B). To validate these results, we performed high resolution scanning electron microscopy (SEM) of beta cells from AL/CR mice after 2 or 12 months on diet to quantify beta cell mature insulin granule and mitochondrial density in individual cells. As expected, we found no differences in beta cell insulin granule content, and we observed higher mitochondrial mass in CR beta cells – which is likely explained by reduced rates of mitophagy (Figure 6B-C, Figure S8C).

**Figure 6.**
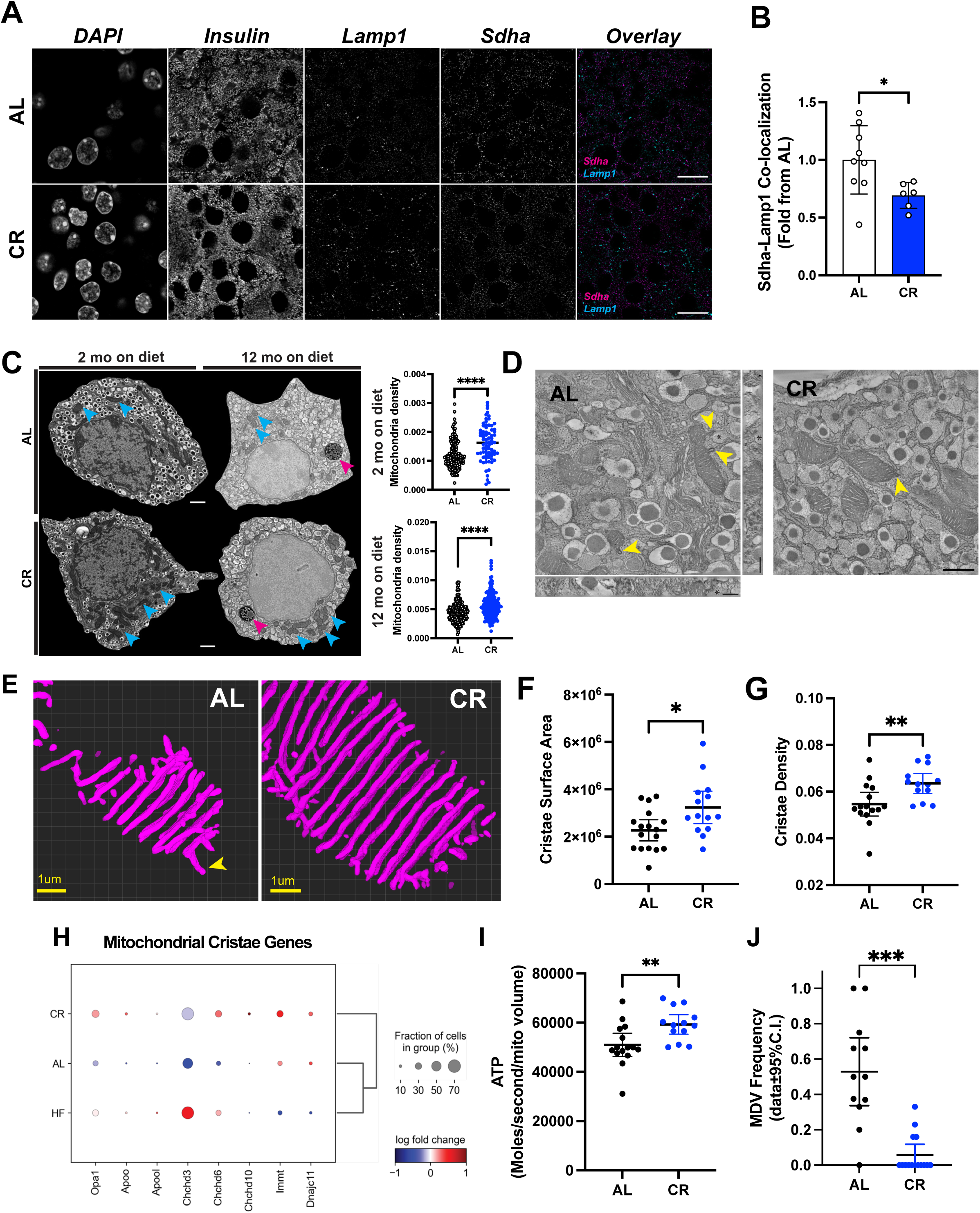
CR increases beta cell mitochondria density and modifies cristae structure. **(A)** Representative imagens of pancreatic sections from AL and CR male mice after 2 months on diet. Slides were stained with Insulin, Lamp1 and Sdha. **(B)** Co-localization between Sdha and Lamp1 was measured. Each dot represents the average co-localization index in each animal calculated from approximately 100 beta cells per animal**. (C)** Representative images of pancreatic beta-cells from AL and CR male mice using electronic microscopy. Blue arrows point to mitochondria density, while pink arrows point to lipofuscin granules in aged beta cells. Mitochondrial density measured by the total mitochondrial number per beta cell area. Each dot represents a single beta cell analyzed from three different islets per mouse (n=3 mice per diet group). **(D)** Representative images of pancreatic beta cells from AL and CR male mice using high resolution electron tomography (eTomo). **(E)** Representative 3D reconstructions of beta cell crista segmentation generated using deep learning image analysis tools. **(F-G)** Cristae surface area and cristae density in AL or CR beta cells. **(H)** Dot plot showing the relative expression levels of genes involved in cristae formation and morphology. **(I)** Calculation of ATP generation for an average beta cell mitochondrion. Each dot represents data from one eTomo image stack. **(J)** Relative frequency of mitochondrial-derived vesicles observed in eTomo images. In (**F-H** and **I-J**), each dot represents an individual field of view with an average of 5 mitochondria per field. eTomo data acquired from n=3 mice per diet group, 5 fields of view per animal. Total of 69-70 mitochondria analyzed per diet group. The asterisks indicate * *p*< 0.05, ** *p*< 0.01, *** *p*< 0.001 and **** *p*< 0.0001 using unpaired Student t-test. In (**D),** scale bar = 500nm, and yellow arrowheads indicating the location of MDVs, whereas the asterisk in each image marks the location of a lysosome.

Given that CR-induced autophagy in beta cells does not target existing insulin granules or mitochondria, and pathways involved in the degradation of beta cell ER (called reticulophagy) are not differentially modulated by CR (Figure S8D), we decided to measure the protein levels of the adaptor protein p62/Sequestosome-1 (*p62/Sqstm1*) since its gene expression levels are increased in CR beta cells (Figure S8B). *p62* targets poly-ubiquitinated (Ub) protein complexes to proteasome and lysosome machineries ^56^. Confocal microscopy of *p62* and *Lamp1* revealed that CR increases beta cell p62-vesicle density without affecting the relative co-localization of p62 with lysosomes, thus indicating that these cells undergo a proportional increase in the number of autophagic events and autophagic flux (Figure S8E). Moreover, CR beta cells have higher expression of ubiquitin processing and Ub protein degradation genes (Figure S5A), including most proteasome subunit genes (Figure S8F), thus indicating that CR-induced autophagy in beta cells likely targets poly-ub macromolecules for degradation via the lysosomal and proteasome machineries.

In addition to reduced mitophagy and higher mitochondrial density, CR beta cells have increased expression of several mitochondrial function genes, including members of the electron transport chain (e.g., *Cox* genes, outer membrane pore complex *Vdac1*)) and higher mitochondrial density (Figure 6 and Figure S8F). We hypothesized that this phenotype would be translated into higher mitochondria efficiency required to maintain beta cell metabolic homeostasis (Figure 4). To investigate beta cell mitochondria structure-function *in situ*, we applied EM tomography (eTomo) using AL and CR mouse pancreases from our 2-month cohort prepared for SEM (see methods for details, Figure S9A)). eTomo provides high-resolution measurements of cellular and mitochondrial structure that can be used to estimate the functional capacity of each mitochondrion to generate ATP and OXPHOS potential from anatomical measurements of mitochondria morphology, cristae number, cristae membrane surface area and density, and type of cristae structure (e.g., lamellar versus tubular)^57^. We reconstructed the 3D architecture of individual beta cell mitochondria and the morphology of mitochondrial cristae with an isotropic axial resolution of 1.62nm using 300 nanometer-thick beta cell sections (Figure 6D, Figure S9A, Video S1), and 3D segmentation of individual cristae was achieved using deep learning-based segmentation pipelines as previously established by us ^58^. This volumetric super resolution EM imaging approach revealed that most beta cell cristae are of the tubular type, and that CR increases beta cell mitochondria cristae surface area and cristae density (Figure 6E-G, Figure S9B-C) without altering mitochondrial volume (Figure S9C). Notably, most beta cell mitochondria contained cristae of the tubular morphology (Figure S9D), which is regulated by proteins in the *Opa1* and MICOS complexes ^59^. In line with our eTomo data, expression of several genes important for cristae structure formation are up regulated in CR beta cells (except for *Apool* and *Chchd3*, Figure 6H). Finally, using all these anatomical measurements of beta cell mitochondria, we calculated the rate of ATP molecules generated per second per mitochondrial volume using an established biophysical mathematical model ^60^ (see methods for details). We estimate that each CR beta cell mitochondria produces ∼59,000 ATP molecules/second/mitochondrial volume, which is 14% higher than in AL beta cells (Figure 6I).

While analyzing the eTomo volumes, we noticed the occurrence of vesicle-like structures stemming from the outer membrane regions of beta cell mitochondria that resemble mitochondrial-derived vesicles (MDVs, Figure 6D, J and Figure S9E-G). MDVs are implicated in mitochondrial proteome quality control pathways by delivering oxidized/damaged macromolecular complexes to the lysosomal degradation system and/or can act as vehicles of inter-organelle transport ^61,62^. We found MDVs in ∼50% of AL beta cell mitochondria, whereas MDVs in CR beta were very rare (Figure 6J). This could be explained by higher expression of oxidative molecular scavenger proteins superoxide dismutase 1 and 2 (*Sod1/2*, Figure S9H). Accordingly, most beta cell MDVs contained membrane-rich cargo that could be seen fusing with lysosomes (Figure S9E-F), while one example fused with the ER. This data suggests that beta cell MDVs are mostly implicated in mitochondrial protein quality control by delivering cargo to the lysosome machinery (Figure S9G).

These results show that CR increases beta cell autophagy to up regulate turnover of macromolecules via the ubiquitin and lysosome systems, while the mitochondrial structure of these cells is protected by a specific reduction in mitophagy. Furthermore, CR-dependent increases in beta cell mitochondria cristae density are expected to facilitate increased ATP production while the reduced formation of MDVs are indicative of reduced accumulation of mitochondrial protein oxidative damage and/or slower rates of mitochondrial proteome turnover.

### CR promotes beta cell longevity

To quantify beta cell longevity, we applied stable isotope labelling of mice using ^15^N-enriched diet followed by a correlated SEM and <u>m</u>ulti-isotope <u>m</u>ass <u>s</u>pectrometry (MIMS) microscopy approach (called MIMS-EM) developed previously by us to quantify cell and protein complex age ^4,7^. Here, Nitrogen 15 (^15^N)-labelled animals were created *in utero* and maintained on ^15^N-diet until post-natal day 45 (P45). Next, ^15^N-mice were randomly allocated to AL, CR, or HFD feeding groups using ^14^N-rich food pellets and maintained on their respective diets for 12 months (i.e., the chase period). Finally, the mice were prepared for MIMS-EM imaging and quantification of cell age based on the amount of ^15^N retained in the cell nucleus (Figure 7A-B) ^4^.

**Figure 7.**
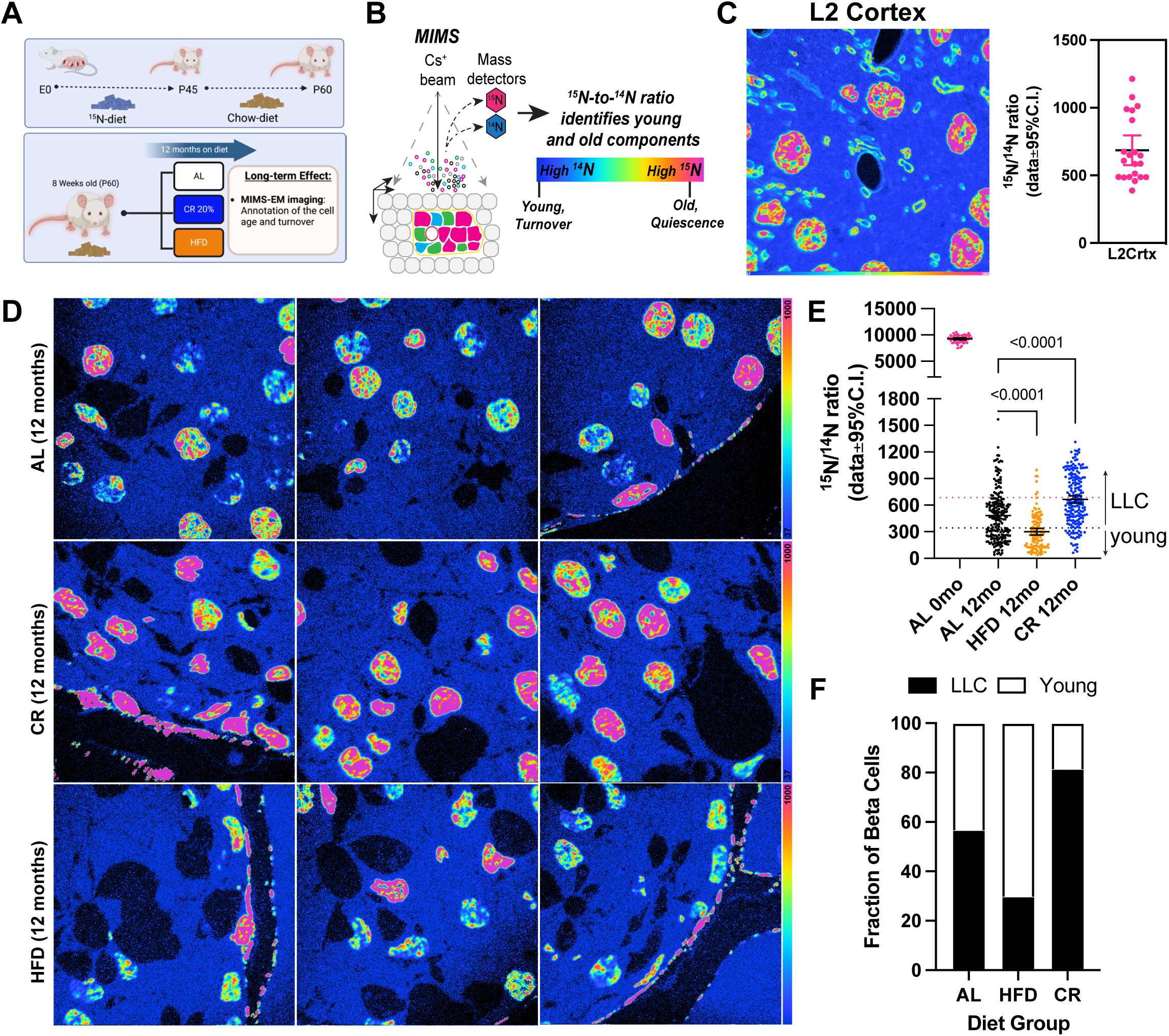
CR promotes beta-cell longevity by slowing-down beta-cell turnover rates. **(A)** Study design. After 2 months of age, ^15^N-labelled mice were kept on AL, 20% CR or HFD for 12 months. **(B)** Multi-isotope imaging mass spectroscopy (MIMS) representation technique. **(C)** MIMS imaging and data quantification of layer 2 cortical neurons from ^15^N labelled mouse after chase with ^14^N diet. Each dot represents a single neuronal nucleus. **(D)** MIMS of pancreatic islets from AL, CR or HFD mice. **(E)** Scatter plot with 15N/14N ratios for beta cells at day 0 (no chase) and after 12 months (mo) on AL, HFD, or CR diet. Each dot represents individual beta cells. The dotted horizontal pink and black lines represent the mean and the lowest ^15^N/14N levels found in cortical neurons shown in (**C). (F)** Estimation of beta cells that are estimated to be LLC or young after 12 months on AL, CR or HFD.

To quantify the maximum amount of ^15^N incorporated by P45 as well as the ^15^N levels remaining in the nucleus of post-mitotic cells after a 1-year lifespan on ^14^N-diet, we performed MIMS-EM of brain neurons after 6 or 12 months of chase (Figure 1C and Figure S10A-B). This approach revealed that P45 mice were saturated with ^15^N and, after 6-12 months of chase most of the ^15^N signal was restricted to the cell nucleus and myelin structures (Figure 7C and Figure S10A-B), as expected ^6,12^. Next, we applied MIMS-EM to islet cells from AL, HFD, or CR mice and found that beta cells at P45 were fully saturated with ^15^N, just like neurons (Figure S10C). Importantly, the levels of nuclear ^15^N in islet and acinar cells were heterogeneous, which is indicative of distinct populations of beta cells with different longevities (Figure 7D) ^4^. We found that HFD beta cells had significantly lower ^15^N levels than AL mice (Figure 7D-E), while CR beta cells had higher ^15^N levels (Figure 7D-E). No effects of HFD or CR on alpha or acinar cells was observed; however, delta cells in CR islets had higher ^15^N levels (Figure S10D). We then classified each beta cell as a potential long-lived cell (LLC) or a relatively younger cell according to their nuclear ^15^N content using cortical neurons as a reference^4^, and found that up to 80% of all beta cells in CR mice are LLCs (Figure 7F). Surprisingly, approximately 30% of all HFD beta cells were LLCs and suggests that a small population of beta cells may be resistant to HFD-induced proliferation (Figure 7F).

## Discussion

In this study we investigated how CR impacts adult beta cell insulin release, metabolism, transcriptional heterogeneity, and aging signatures. We exposed adult male and female mice to a 20% reduction in daily caloric intake by providing a limited amount of food before the start of their active period at night. Recent studies have shown that this level of CR is associated with long fasting periods that are important for the beneficial metabolic and geroprotective effects of CR, and which are mostly observed in male mice ^29,63,64^. Accordingly, 20% CR prevented the accumulation of fat mass while preserving lean mass in both sexes; however, only male mice showed improvements in glucose homeostasis and higher peripheral insulin sensitivity, thus highlight existing sex-dependent differences in glucose homeostasis mechanisms during CR and diabetes ^21,65,66^.

Increased insulin sensitivity and/or enhanced insulin signaling during CR are linked to increased longevity in several organisms, from worms to humans ^67–69^. In contrast, insulin resistance and/or hyperinsulinemia occur during type 2 diabetes (T2D) and are implicated as the main causes of T2D ^70^. CR has been proposed as a feasible approach to combat cardio-metabolic diseases like T2D due to its positive effects on whole body insulin sensitivity that reduce the metabolic demand for insulin release by lowering blood glucose levels. However, here we show that lower insulin release levels occur independently of changes to blood glucose levels. This reduced insulin secretion tone is adequate to maintain euglycemia without triggering hypoglycemic events after meals (even after exposure to incretin receptor agonists). This means that CR promotes a physiological scenario where beta cells “rest” due to a significantly lower metabolic demand for insulin secretion. A similar strategy has been proposed for treatment of T1D and T2D, where exogenous insulin and/or the KATP channel opener diazoxide are applied to recover beta cell function (at least temporarily) ^71,72^. To understand how a lower metabolic demand affects the heterogeneity of beta cells during CR, we used single nucleus transcriptomics and found that CR promotes activation of beta cell function, identity, and protein and organelle homeostasis pathways. By profiling beta cell chromatin accessibility and TF activity, we show that CR beta cells activate genes with TF motifs targeted by modulators of circadian (*Dbp*, *Bmal*), beta cell identity (*Mafa*), and ER stress and cell homeostasis (*Atf6*, *Creb3l2*) pathways. These findings are in line with previous studies showing that CR achieved via time-restricted corrects abnormal glucose homeostasis by activating *Dbp*^33^, and that the circadian rhythm regulator *Bmal* is important for maintenance of mature beta cell identity and function^73^.

We validated our transcriptomics using light, electron, and mass spectrometry microscopy; first, we found higher *Bmal* and *Pdx1* expression as well as increased density of autophagy vesicles marked by *p62/Sqstm1*, *Lc3I-II*, and *Lamp1* in CR beta cells *in situ*. Previous studies have shown that CR and CR via TRF restore beta cell function in HFD-fed animals by rescuing gap-junction connectivity and autophagic fluxes ^30,34^. In this context, beta cell autophagy is implicated in the degradation of insulin granules and/or mitochondria to protect beta cells from accumulating molecules and/or damaged organelles during pro-inflammatory and metabolic stress conditions ^12,34,74–77^. In contrast, our studies reveal that CR-induced autophagy in beta cells does not target insulin granules nor the mitochondria; instead, autophagy is involved in the degradation of poly-ubiquitinated substrates delivered to proteasomes and/or lysosomes via *p62*-positive autophagy vesicles. Importantly, CR prevented the loss of autophagy vesicles during aging, and reduced beta cell senescence signatures^19,53^, as evidenced by a reduction in the accumulation of DNA damage and in the expression of beta cell senescence markers *p21/Cdkn1a* and *p16/Cdkn2a*, and in the relative increase in nuclear *LmnB*1 levels.

The reduced levels of mitophagy observed in CR beta cells are expected to contribute (at least in part) to higher mitochondrial density, while the elevated expression of multiple electron transport chain, cellular respiration, and cristae morphology genes suggest that mitochondrial metabolism and homeostasis are optimized in CR beta cells. This hypothesis is supported by three separate lines of observations. First, our MALDI MS metabolomics experiments reveal normal-to-elevated levels of lipids, cAMP, glucose metabolites, and ADP-ribose – the later which could explain why CR beta cells accumulate less DNA damage^51^, which in turn explains the reduced expression of senescence markers ^18,19,52^. Second, using eTomo microscopy, we discovered that CR increases mitochondrial cristae density that translates into a higher potential of each mitochondrion to generate more ATP. Given that insulin release is reduced in CR cells, this increased ATP production may serve as fuel to sustain ATP-demanding processes activated during CR such as protein ubiquitination and degradation in proteasomes, and autophagy^78^. Third, the near absence of MDVs in CR beta cells and up regulation of superoxide scavengers *Sod1* and *Sod2* strongly suggests ROS generation and degradation of mitochondrial proteins is reduced during CR. Furthermore, previous studies show that CR increases mitochondrial bioenergetic efficiency and reduces oxidative stress accumulation in different systems^21,79^ (including in humans^80^), which is linked to increased mitochondrial protein longevity in aged cells^81^. Reduced accumulation of oxidative damage (in the form of reactive oxygen species (ROS)) and enhanced bioenergetics of CR beta cells could also explain why these cells secrete less insulin during fasting, as dysregulation of ROS and glycolytic signaling are proposed as causes of basal hyperinsulinemia during diabetes ^70,82^. Future experiments will focus on measuring mitochondrial energetics under different basal and stimulated conditions to further understand how CR regulates basal insulin secretion while promoting healthier mitochondria homeostasis in beta cells.

Finally, using ^15^N-SILAM and MIMS-EM we have previously shown that most mouse beta cells are long-lived while the rest constitutes a sub-population of beta cells that proliferates throughout their lifetime ^4^. Using this same approach, we show that CR prolongs the longevity of beta cells by reducing the turnover rate of the subpopulation of beta cells that is more proliferation prone, while HFD increases beta cell proliferation (as expected). Strikingly, we also found long-lived beta cells in the HFD setting, thus indicating that some beta cells do not proliferate during long periods of hyperglycemia *in vivo*. Previous studies have shown that proliferation promotes an immature and less functional beta cell phenotype than non-proliferating beta cells ^83^, which is also supported by the higher levels of beta cell identity and function markers in CR islets. Importantly, this largely post-mitotic state of CR beta cells could also explain the reduced accumulation of DNA damage, as cell proliferation triggers higher accumulation of genetic mutations and replication-dependent stress that can lead to disease and aging ^84^.

In conclusion, CR enhances beta cell health and longevity by upregulating cell identity and energy homeostasis mechanisms while delaying beta cell aging. Therefore, these results offer a molecular footprint of how CR affects beta cells to preserve and/or rescue beta cell dysfunction during aging and T2D.

### Limitations of our study

Our report shows how CR impacts mouse beta cell heterogeneity, including their in vivo function and molecular and aging signatures. In our experimental setup, CR was efficiently achieved by providing a restricted amount of food before the start of the animal’s active phase. By design, this approach potentializes the metabolic impacts of CR (including the periods of prolonged fasting) by activating circadian rhythm mechanisms involved in nutrient metabolism ^29,65^. Therefore, we cannot distinguish between the effects of recurrent and prolonged fasting periods versus reduced daily calorie intake (without fasting) on beta cell function and heterogeneity. Importantly, recent studies show that combining CR with fasting leads to better metabolic and antiaging effects ^29,63,65^. Our single cell transcriptomics and GRN reconstructions identify likely regulators of the beta cell phenotype described here. Therefore, future experiments will focus on exposing genetic mouse models (i.e., *Mafa* beta cell knockouts) to CR. The ^15^N-SILAM and MIMSEM experiments add to previous observations of cellular age mosaicism in somatic tissues, including in the pancreas ^4,7,8,84^. Like MALDI MS metabolomics and eTomo, this approach is time consuming and financially expensive (thus limiting the number of animals and cells analyzed).

Notably, the nuclear ^15^N measurements of any cell are variable since heterochromatin-rich domains in the sections analyzed are randomly sampled during MIMS-EM. These compartments tend to have higher density of ^15^N-DNA and yield higher signal intensities versus DNA “poor” euchromatin domains ^4^. Nevertheless, our results confirm previous MIMS-EM results, and now show how dietary interventions such as CR can be used to promote a long-lived beta cell phenotype.

## Acknowledgements

This work is dedicated to Professor Claude P. Lechene, who sadly passed away in 2023. We are thankful to Claude for his ingenuity and dedication to the stable isotope labelling and imaging fields, and for his generosity in sharing with us his time and knowledge that has led to the development of MIMS-EM and other multi-scale imaging modalities. We are also thankful to Michelle Reyzer from the Vanderbilt Mass Spectrometry Research Center for assistance with MALDI MS experiments. This research was supported by recruitment funds from the Vanderbilt’s Department of Molecular Physiology and Biophysics, a HIRN Gateway (R03 DK127484) and a DKNET (5U24DK097771) awards to RAeD, by a grant from the Canadian Institutes of Health Research (CIHR; 487188) to PEM. PEM holds the Canada Research Chair in Islet Biology. This research was also supported by NIDDK 5R01DK132669-02 to DD, and by NIH grants R24GM137200, U24NS120055, 1S10OD021784, and R01 DA038896, as well as the National Science Foundation (NSF) grant NSF2014862-UTA20-000890 to MHE.

## Data Availability

Data and code used in this manuscript is available upon request and deposited here: DOI: 10.17632/wskdgg72k2.1

## Author Contributions

CDS, MH, MHE, and RAeD designed experiments; CDS, SS, MC, GP, VLR, CA, BR, KK, TD, MC, DD, PEM, MH, MHE, and RAeD collected and analyzed data. CDS and RAeD wrote the manuscript with significant input from all authors.

## Declaration of Interest

The authors declare no competing interests.

---

**Supplementary Figure 1. (A)** Cumulative energy intake over 2 months on diet from FVB male mice fed with AL, 20% CR or HFD. Each point represents the average from each cage with five mice, n=4-10 cages per experimental group from two different sets. **(B)** Percentage of fat and lean mass per body weight from AL and CR male mice. **(C)** Blood glucose levels during the oral glucose tolerance test (oGTT) after 2 months on diet and respective area-under-curve (AUC) measurements. **(D)** Insulin levels during the oGTT and **(E)** the respective ratio between the insulin and glucose values. **(F)** Insulin degrading enzyme activity measured in liver samples from male mice after 2 months on diet. **(G)** Pancreatic islet morphology from male mice after 2 or 12 months on diet. Beta, alpha and delta cells were detected by immunofluorescence, using antiinsulin (yellow), anti-glucagon (magenta) and anti-somatostatin antibodies (white). Quantitative analysis of beta-cell mass and the percentage of beta, alpha and delta-cells per islet were performed in two different sections from each pancreas. **(H)** Dynamic insulin secretion obtained from perifused islets with insulin secretagogues (5.6 mM and 16.7 mM glucose, IBMX 100 mM, and KCl 20mM) and the respective **(I)** insulin total content normalized by IEQ. **(J)** Schematic diagram of mice subjected to liraglutide (400 ug/Kg i.p) 2 hours before the MTT. **(K)** Blood glucose levels during the MTT using mice pretreated with liraglutide and respective AUC measurements. **(L)** Insulin levels during the MTT and **(M)** the respective ratio between the insulin and glucose values. Data are presented as mean ± 95% Cl. The asterisks mean * *p*< 0.05, ** *p*< 0.01, *** *p*< 0.001 and **** *p*< 0.0001 using two-way ANOVA with Tukey’s post hoc test when three groups are compared or unpaired Student t-test for two groups comparation. In **(B)**, n=7 male mice per diet group; **(C-E)** n=5 male mice per diet group; **(F)** n=6-9 male mice per diet group; **(G)** n=5 male mice per diet group; **(H-I)** n=5-10 male mice per diet group. **(K-M)** n=8 male mice per diet group.

**Supplementary Figure 2. (A)** Daily body mass of C57BL6J male mice fed with AL or 20% CR for 2 months. **(B)** Cumulative energy intake over 2 months on diet, each point represents the average from each cage with five mice. **(C)** Body mass change and **(D)** the ratio of fat and lean mass. **(E)** Blood glucose levels obtained during the meal tolerance test (MTT) after 2 months on diet and the respective AUC. **(F)** Insulin levels obtained during the MTT and **(G)** the respective fold change from baseline values. **(H)** HOMA-IR calculated from fasting glucose and insulin values after 2 months on diet. Data are presented as mean ± 95% Cl. The asterisks mean * *p*< 0.05, ** *p*< 0.01, *** *p*< 0.001 and **** *p*< 0.0001 using two-way ANOVA with Tukey’s post hoc test or unpaired Student t-test for two groups comparation, n=5 male mice per diet group.

**Supplementary Figure 3. (A)** Daily body mass of female mice fed with AL or 20% CR for 2 months. **(B)** Cumulative energy intake over 2 months on diet, each point represents the average from each cage with five mice, n=2-1 cages per experimental group. **(C)** Body mass change and **(D)** the ratio of fat and lean mass. **(E)** Blood glucose levels obtained during the meal tolerance test (MTT) and the respective AUC. **(G)** Insulin levels obtained during the MTT and **(H)** the respective fold change from baseline values. **(I)** HOMA-IR calculated from fasting glucose and insulin values after 2 months on diet. **(J)** Blood glucose levels obtained during the insulin tolerance test (ipITT) and **(K)** the respective decay of the glucose rate per minute (kITT). **(L)** Differences in kITT between male and female mice subject to AL or 20% CR for 2 months. Data are presented as mean ± 95% Cl. The asterisks mean * *p*< 0.05, ** *p*< 0.01, *** *p*< 0.001 and **** *p*< 0.0001 using two-way ANOVA with Tukey’s post hoc test or unpaired Student t-test for two groups comparation. NS means not significant, n=5 female mice per diet group.

**Supplementary Figure 4. (A)** Fraction of mitochondrial (mt) genes expressed in all sequenced cells from AL, CR, and HFD mice. **(B)** Graph with number of genes (features) identified as a function of the number of RNA molecules detected (nCount) in cells from AL, CR, and HFD mice. **(C)** ATAC fragment length in all sequenced cells from AL, CR, and HFD mice. **(D)** Transcription start-site (TSS) enrichment in AL, CR, and HFD samples. **(E)** Pie chart showing the breakdown of ATAC peak distribution in types of genomic loci. **(F)** Graph with the number of peaks and RNA molecules with unique molecular identifiers (UMI) in AL, CR, and HFD cells. **(G)** Representative immunohistochemistry (IHC) and confocal microscopy of AL and CR islets after 2 months on diet using anti-Pdx1 and anti-Bmal antibodies. Graph on the right shows the relative frequency of beta cells with high versus low levels of nuclear *Bmal*. **(H-I)**, differentially expressed genes and chromatin accessibility sites in beta cell subpopulations, respectively, of AL, CR, and HFD islets. **(J)** List of genes mapped closest to enriched accessible chromatin sites in different beta cell populations.

**Supplementary Figure 5. (A-E)** Dot plot with hierarchical clustering analysis (HCA) highlighting the expression levels of genes associated with **(A)** ion channels, insulin secretion, vesicle organization, protein folding, ubiquitin processing and endoplasmic reticulum (ER) stress in each betacell state; **(B)** autophagy, amino acid activation and DNA damage in each beta-cell state; **(C)** lipid metabolism in each beta-cell state; **(D)** glycolysis, pyruvate metabolism, tricarboxylic acid cycle (TCA) and pentose phosphate pathway, and **(E)** amino acid transport in each beta-cell state.

**Supplementary Figure 6. (A)** Graph with the number of transcription factors (TFs) identified with SCENIC by the number of targets per TF. **(B)** SCENIC regulon specificity score (RSS) for individual TFs in AL, CR, or HFD beta cells. **(C)** RSS and Z-scores for TFs enriched in AL, CR, and HFD beta cells. **(D)** Pearson correlation matrix of TFs identified in mouse beta cells from AL, CR, or HFD mice. Boxes highlight clusters of TFs with high degree of correlation. **(E)** Venn diagrams illustrating the degree of overlap for *Mafa*, *Foxp1*, or *Creb3l2* regulons.

**Supplementary Figure 7. (A)** Relative pancreatic weight in AL and 20% CR mice after 2 or 12 months on diet. **(B)** Representative images of pancreatic islets stained with Lamin B1 from AL and CR male mice after 12 months on diet and the respective quantification of Lamin B1 high cells per mice. **(C**) Quantification of p21 expression *in situ* in pancreatic beta-cells from AL and CR mice after 12 months on diet. **(D)** XY plot showing the correlation between the number of p16 and p21 mRNA spot-object detected in each beta cell from mice kept on AL or CR for 12 months. Solid and dashed lines indicate the linear regression and confidence intervals, respectively. r2 values are shown on top of each graph. **(F)** Slides were stained with Insulin and phospho-S6 ribosomal protein. **(G)** Quantification of p-S6 intensity in pancreatic beta-cells. Data are presented as mean ± 95% Cl. Data are presented as mean ± 95% Cl. The asterisks mean **** *p*< 0.0001 using one-way ANOVA with Tukey’s post hoc test or unpaired Student t-test for two groups comparation, n=5-15 pancreas per diet group.

**Supplementary Figure 8. (A)** Co-localization analysis between Insulin and the autophagy marker Lamp1 using pancreatic sections from AL and CR male mice after 2 months on diet. Representative images of islets stained with Insulin and Lamp1 are shown in left. **(B)** Dot plot with hierarchical clustering analysis (HCA) highlighting the expression levels of genes associated with mitophagy. **(C)** Quantification of insulin granules per beta-cells using SEM imaging. **(D)** Same as in **(B)** showing genes involved in the reticulophagy pathway in beta-cells from AL, CR and HFD mice. **(E)** Colocalization of beta cell p62 vesicles with Lamp1 and quantification of beta cell area occupied by p62 in AL or CR mice. **(F-G)** Same as in **(B)** showing genes involved in the proteasome and respirasome pathways, respectively, of AL, CR, or HFD beta cells.

**Supplementary Figure 9. (A)** Schematic diagram of eTomo microscopy. **(B)** Average mitochondria cristae density and cristae surface area for each animal analyzed in AL and CR diet groups. **(C)** Mitochondrial surface area to volume ratios in eTomo images. Each dot represents individual mitochondrial objects analyzed. **(D)** Fraction of mitochondria with tubular versus lamellar cristae architecture. **(E-F)** Consecutive z sections showing fusion of a beta cell MDVs with a lysosome. Yellow arrowheads indicate MDVs with multi-membranous structures budding of from the mitochondria outer membrane space and the asterisk marks the lysosome. **(G)** Snapshot of a beta cell MDVs fusing with the ER. In (E-G), yellow arrowheads indicate MDVs locations. **(H)** Dot plot with genes involved in the reactive oxygen species (ROS) pathway in AL, CR, or HFD beta cells.

**Supplementary Figure 10. (A)** MIMS imaging and ^15^N/^14^N ratiometric maps of granule neurons in the cerebellum of ^15^N-labelled mice at day 0 or after 6 or 12 months of chase in AL, HFD, or CR diet groups. **(B)** ^15^N/^14^N values from granule neurons from mice kept on AL, HFD, or CR for 6 or 12 months. Each dot represents a single nucleus. **(C)** SEM and MIMS imaging with ^15^N/^14^N ratiometric maps of islet cells in the pancreas from a ^15^N-labelled mouse at day 0. **(D)** ^15^N/^14^N values in the nucleus of alpha cells, delta cells, and acinar cells from the pancreas of mice kept on AL, HFD, or CR for 12 months. Each dot represents a single cell nucleus. P values are shown.

## Legend for Supplementary Tables and Video.

**Supplementary Table 1.** List of differentially expressed of genes (DEG) in AL, CR, or HFD beta cells.

**Supplementary Table 2.** List of differentially regulated transcription factor motifs in AL, CR, or HFD beta cells.

**Supplementary Table 3.** List of differentially expressed of genes (DEG) in the identified beta cell states of AL, CR, or HFD beta cells.

**Supplementary Table 4.** List of transcription factors identified using SCENIC analysis of AL, CR, or HFD beta cell transcriptomics.

**Supplementary Table 5.** Detailed list of genes associated with regulons for Mafa, Foxp1, or Crebl2 in AL, HFD, and CR beta cells.

**Supplementary Table 6.** Full list of genes associated with all identified regulons in AL, HFD, and CR beta cells.

**Supplementary Table 7.** ROC analysis of metabolite ions identified in islet or acinar compartments of AL or CR pancreases.

**Supplementary Table 8.** Metabolite identity HMDB hit list of top metabolites enriched in AL islets.

**Supplementary Table 9.** Metabolite identity HMDB hit list of top metabolites enriched in CR islets.

**Supplementary Video 1.** Movie of a representative beta cell eTomo acquisition field of view from an AL mouse islet.

## Notes

### Competing Interest Statement

The authors have declared no competing interest.

