## Supplementary figures and images for "Caloric restriction promotes beta cell longevity and delays aging and senescence by enhancing cell identity and homeostasis mechanisms"

### Supplementary Figures

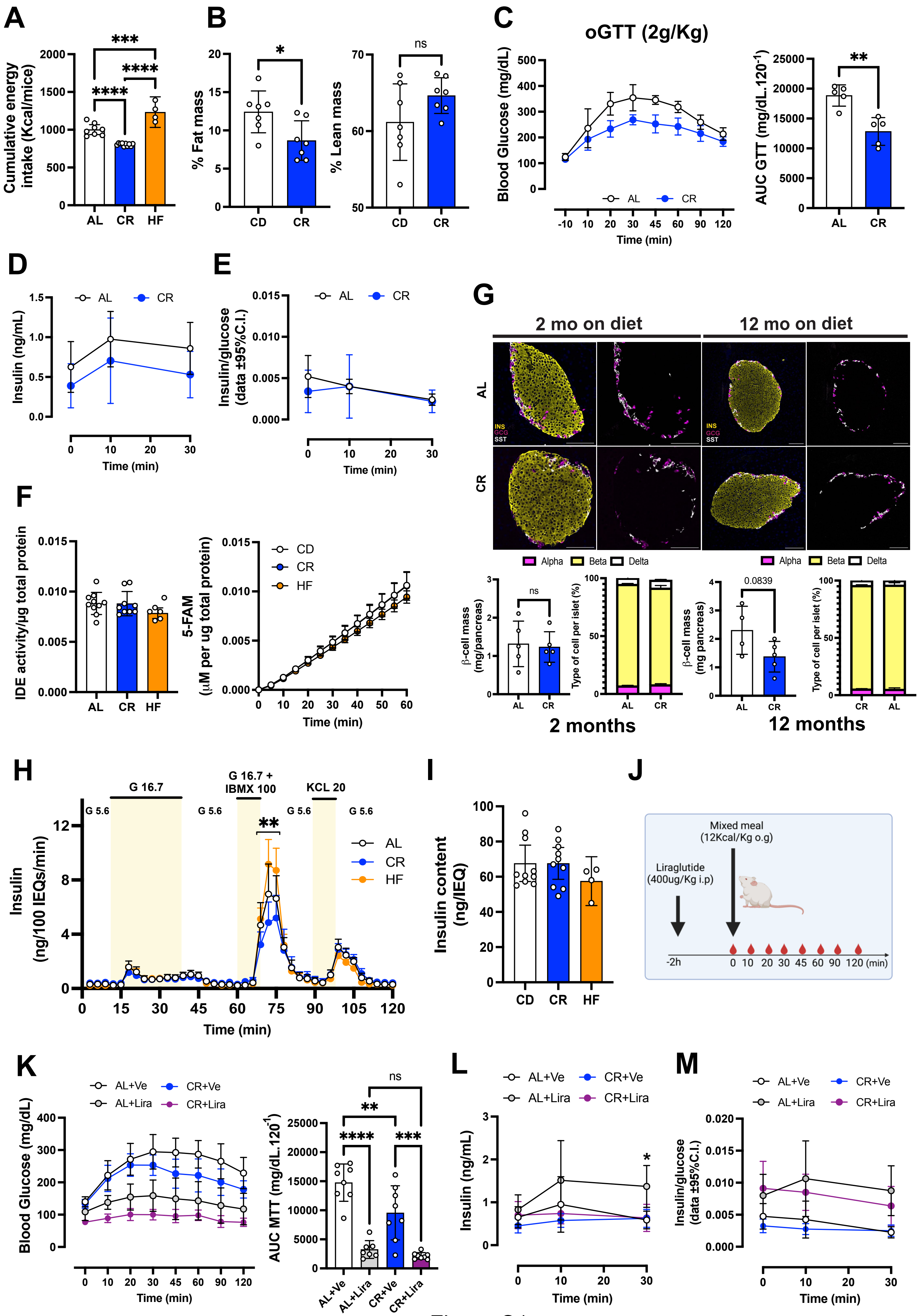

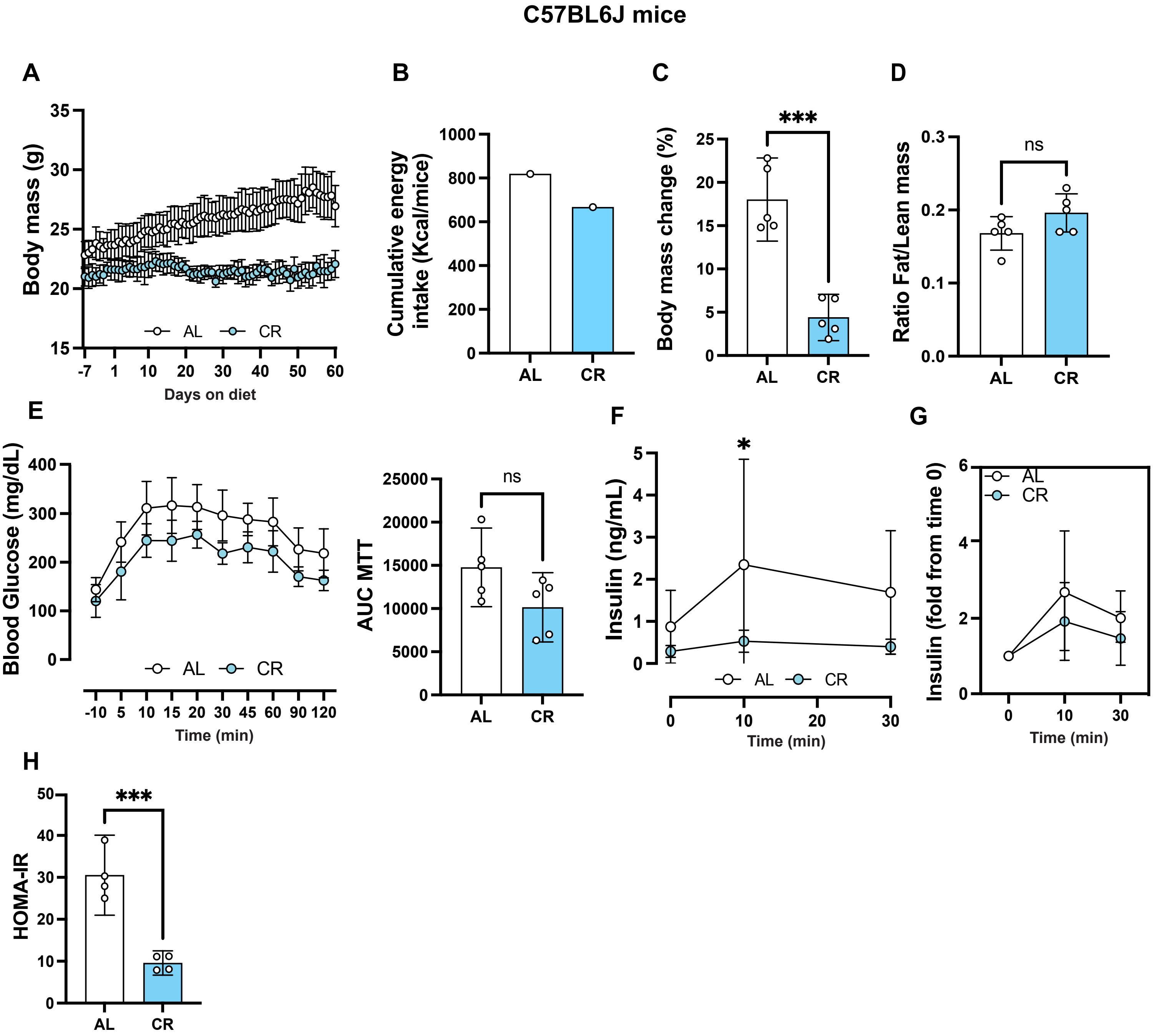

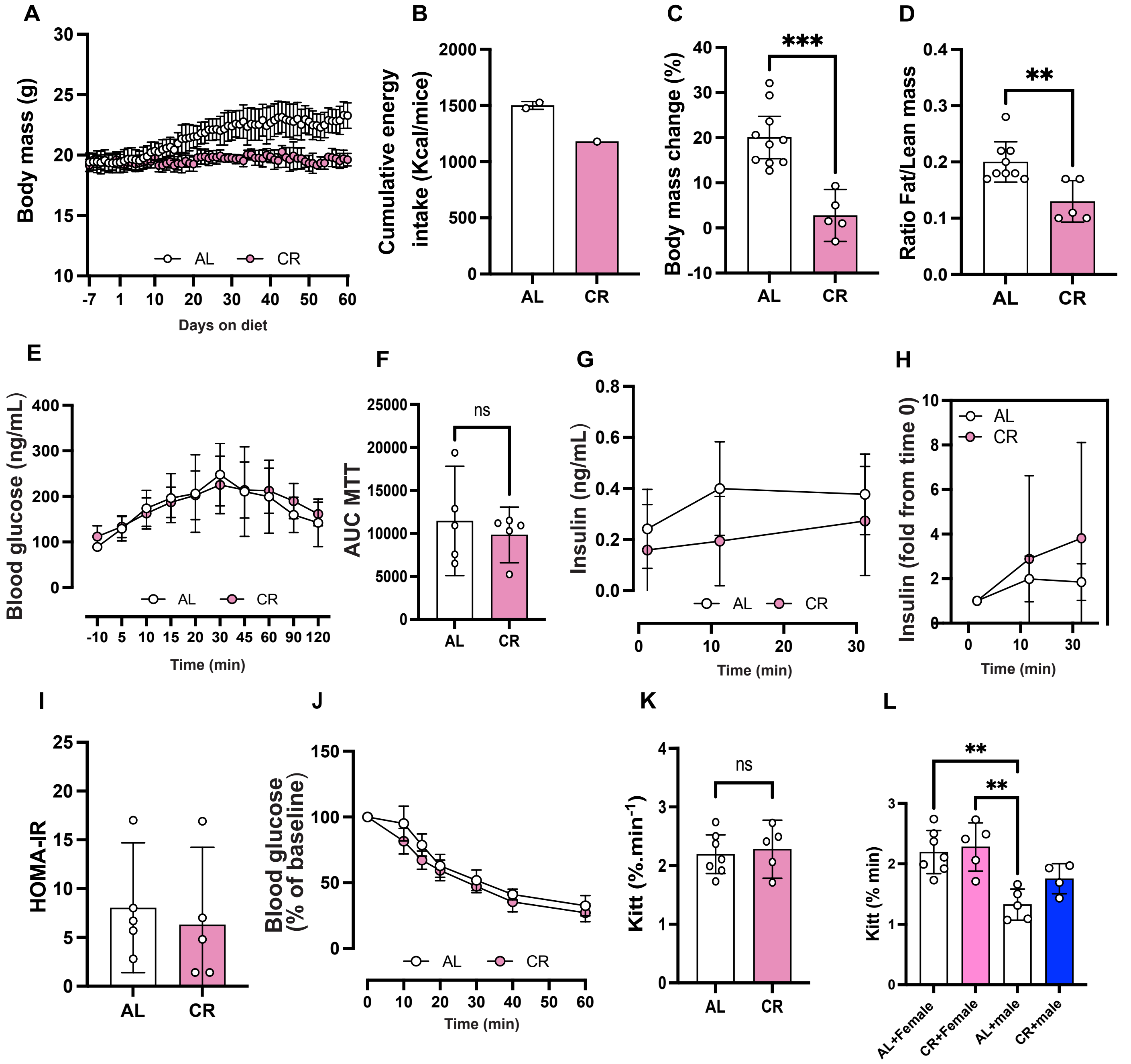

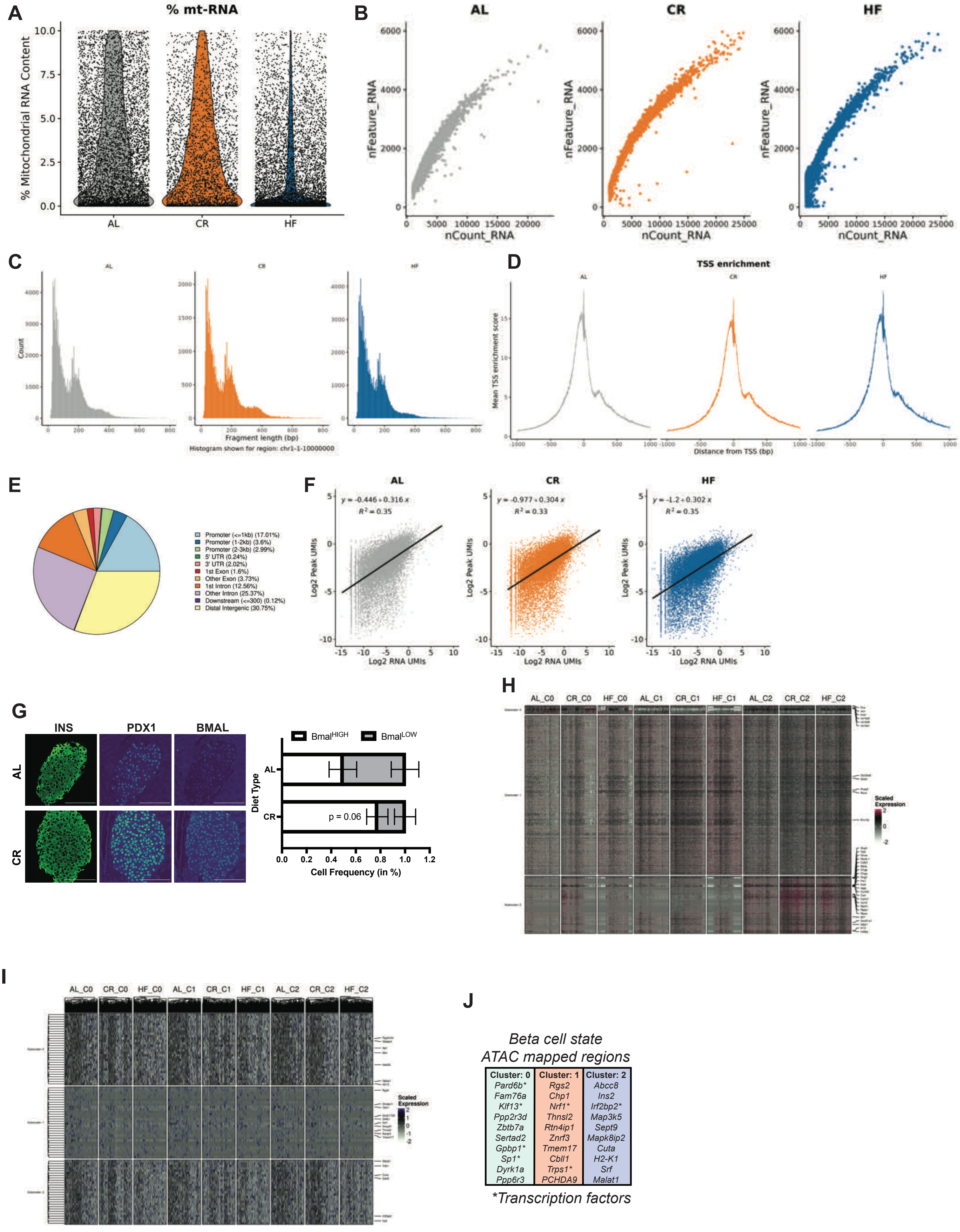

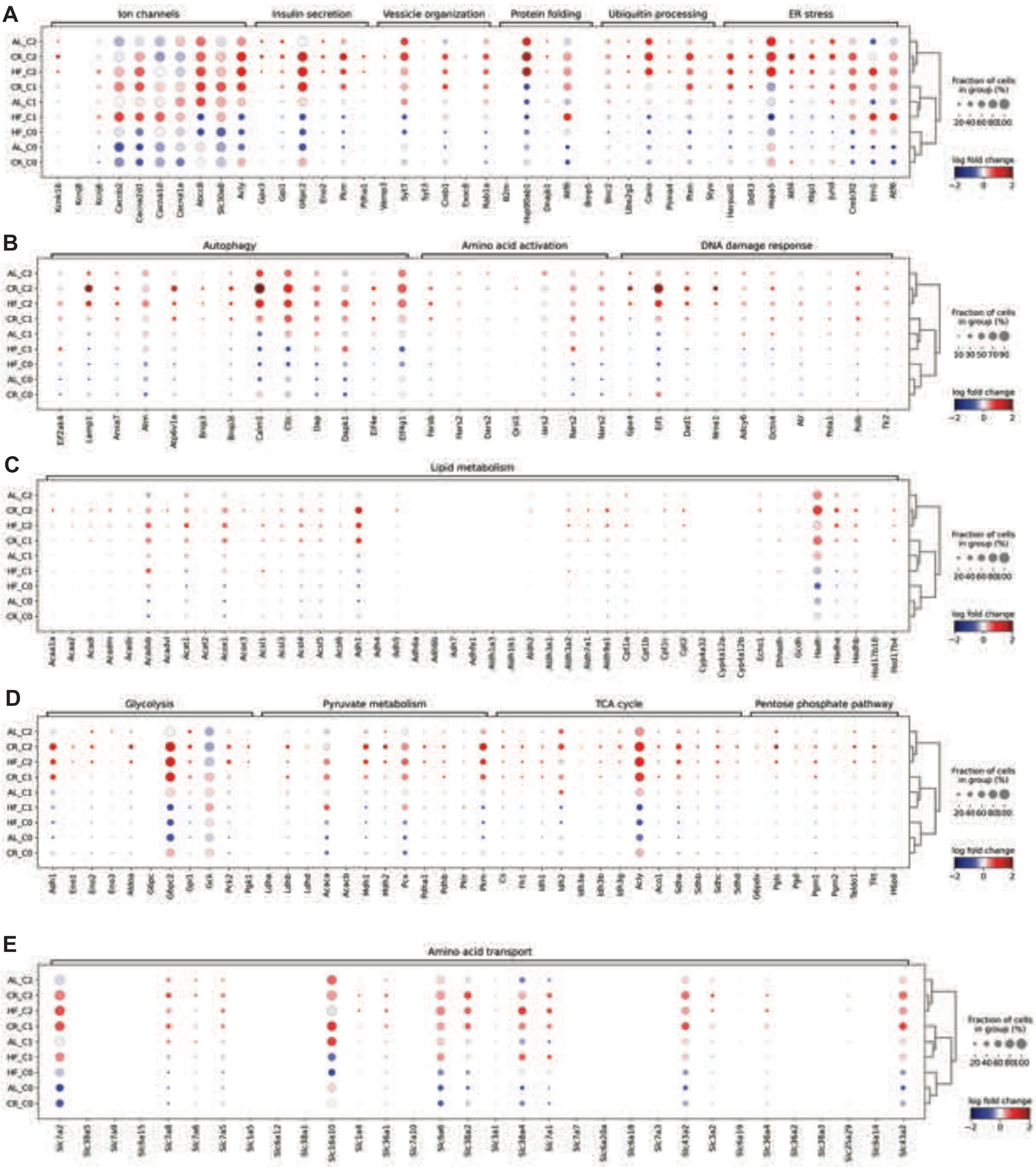

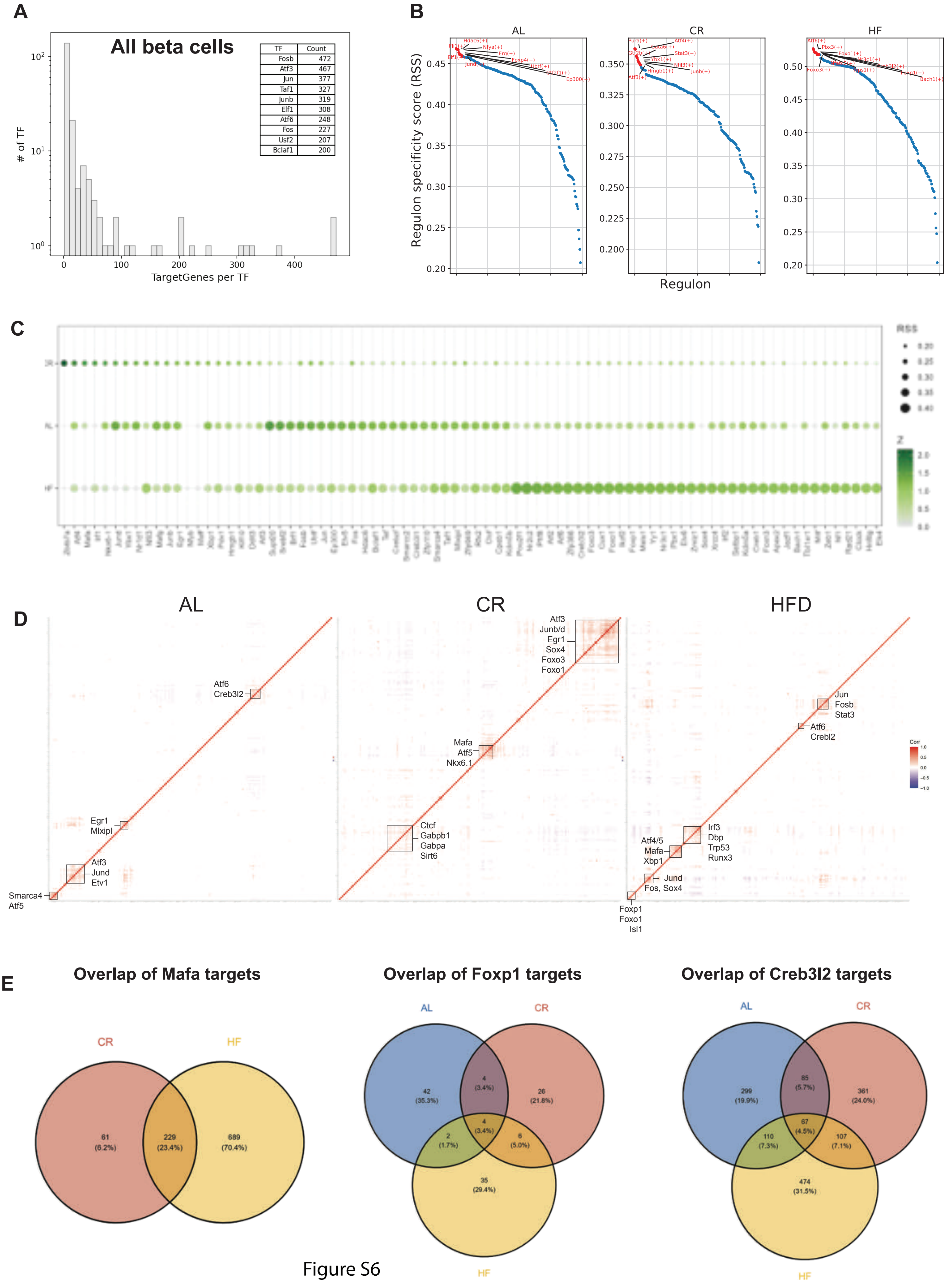

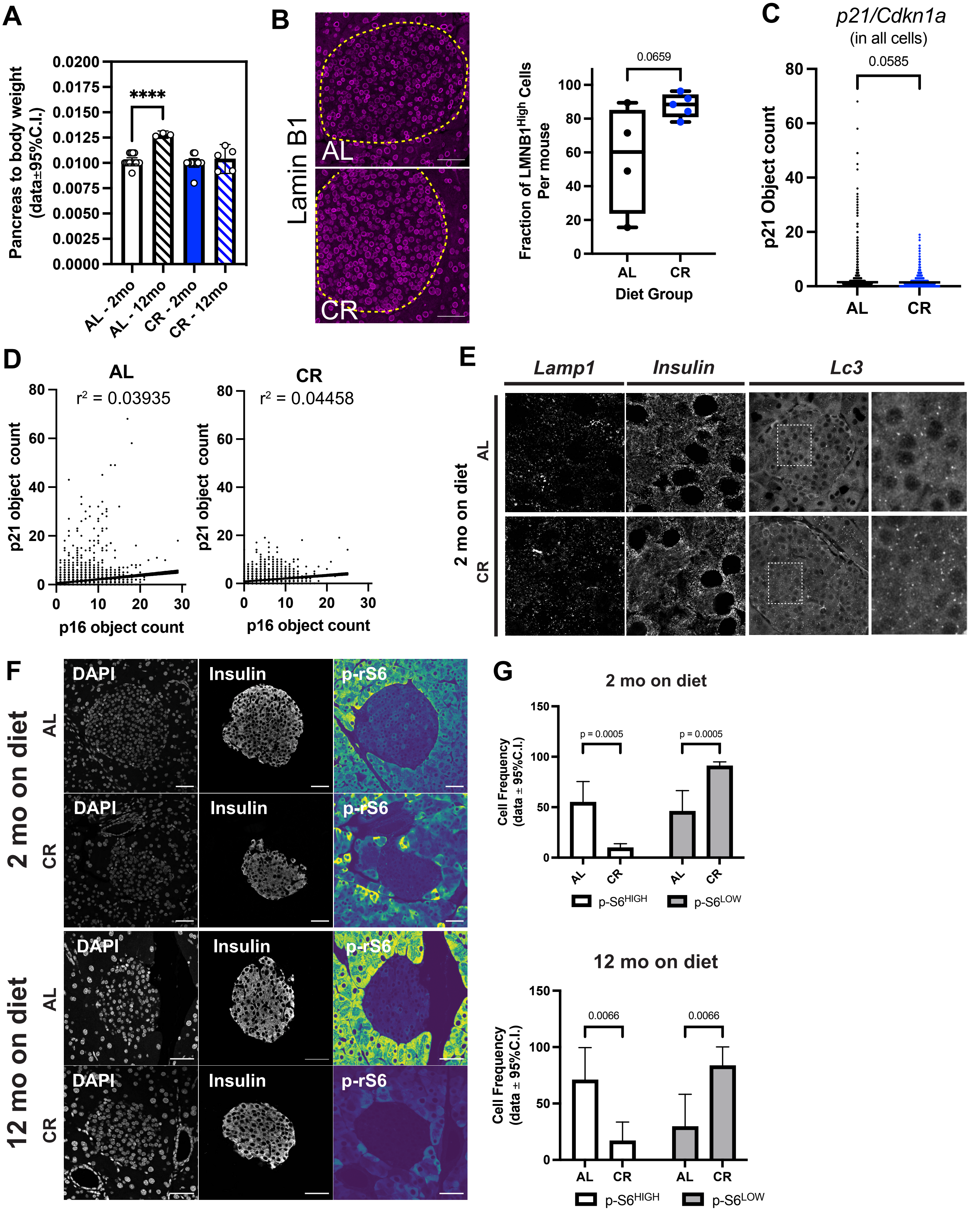

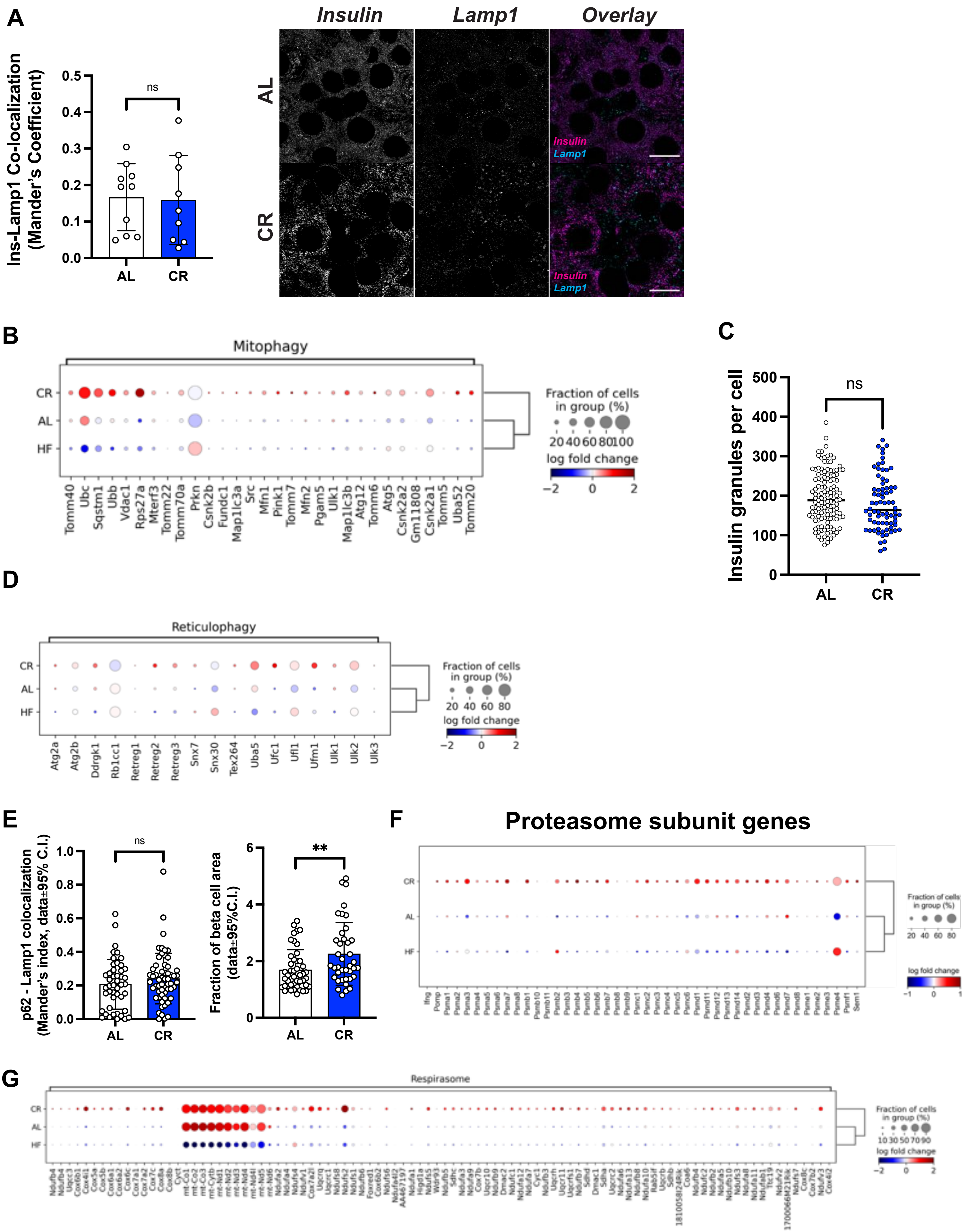

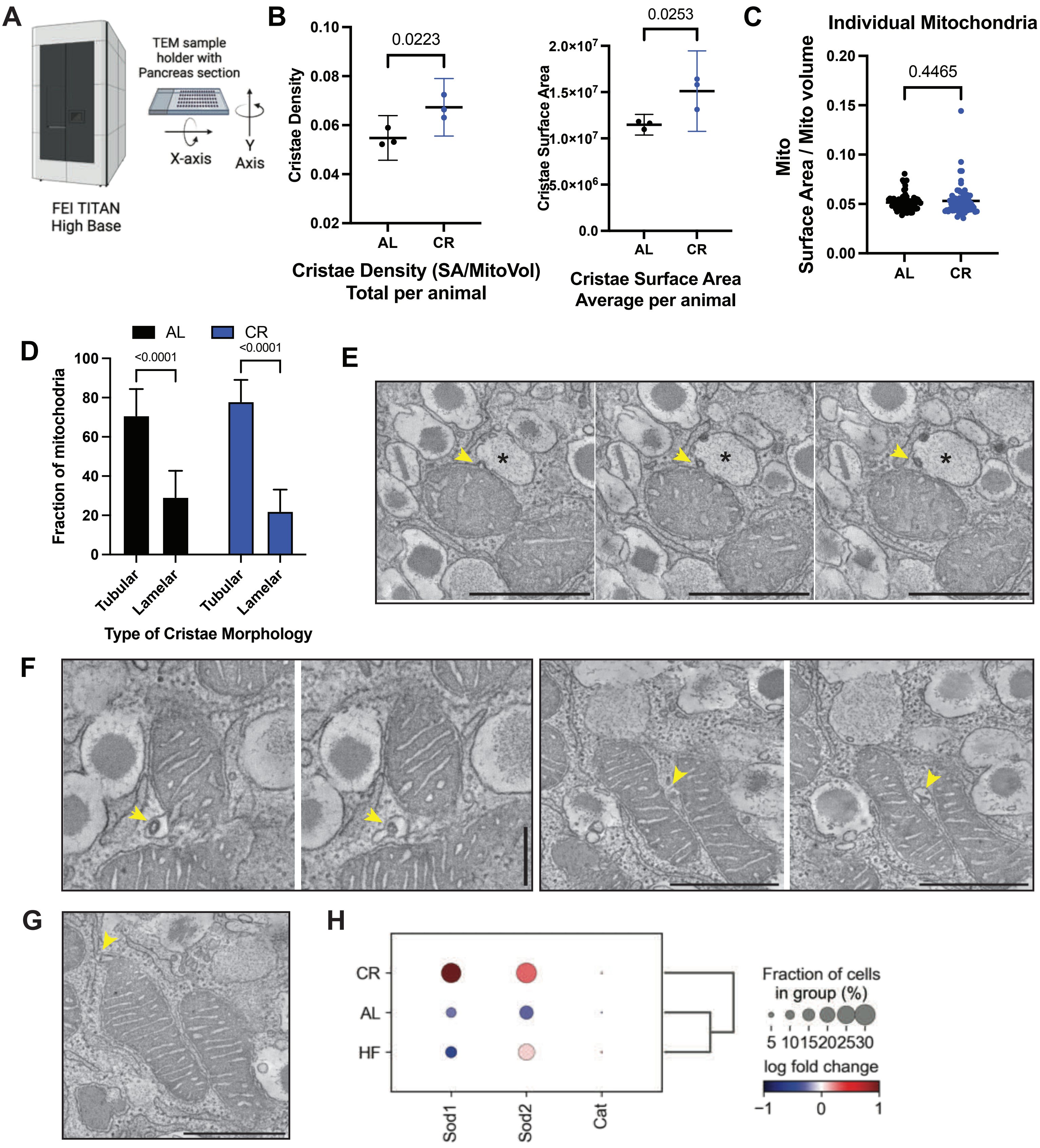

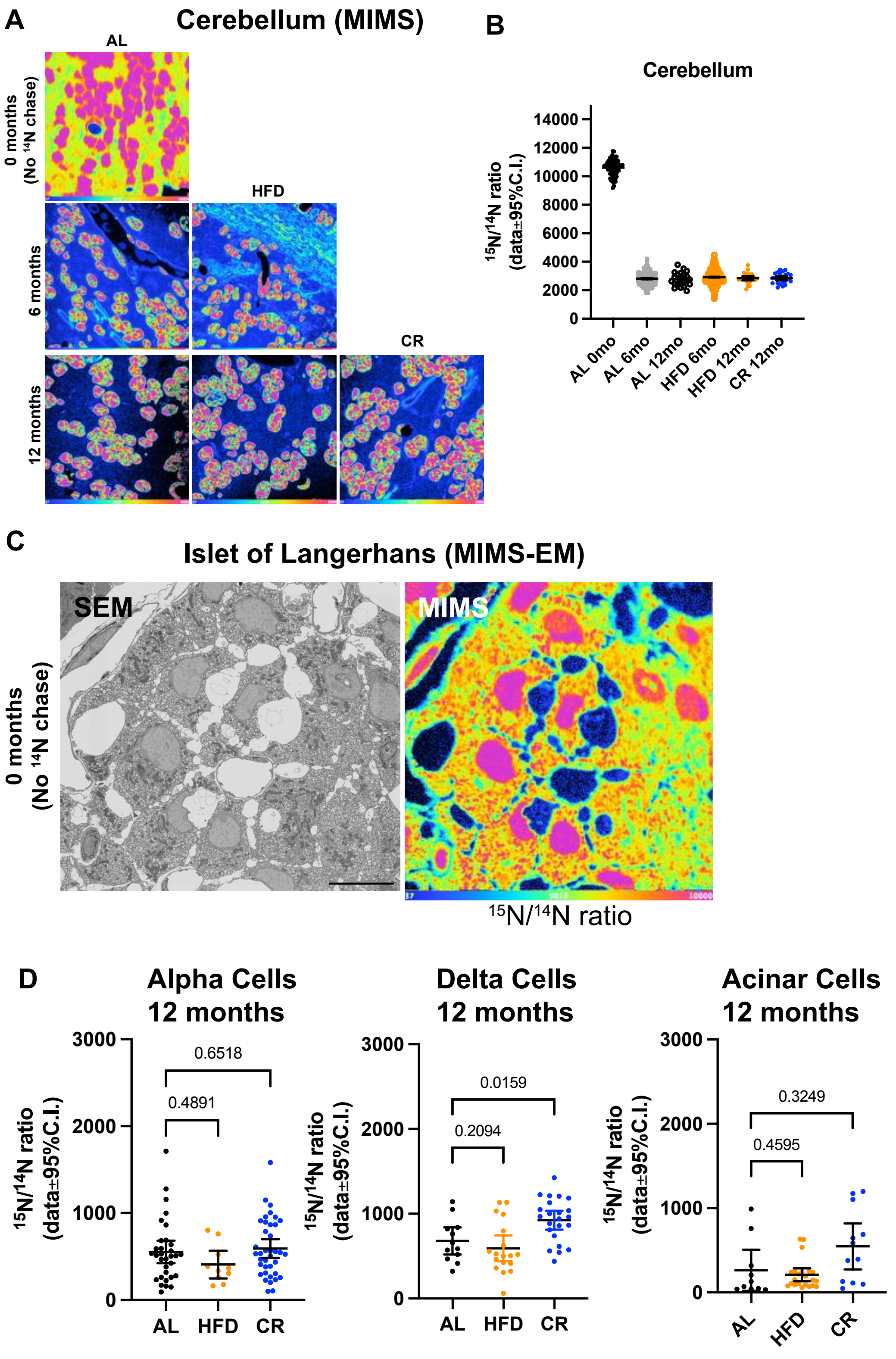
